# Disobind: a sequence-based, partner-dependent contact map and interface residue predictor for intrinsically disordered regions

**DOI:** 10.1101/2024.12.19.629373

**Authors:** Kartik Majila, Varun Ullanat, Shruthi Viswanath

## Abstract

Intrinsically disordered proteins or regions (IDPs/IDRs) adopt diverse binding modes with different partners, from coupled-folding-and-binding, to fuzzy binding, to fully-disordered binding. Characterizing IDR interfaces is challenging experimentally and computationally. The state-of-the-art AlphaFold-multimer and AlphaFold3 can be used to predict IDR binding sites, although they are less accurate at their benchmarked confidence cutoffs. Here, we developed Disobind, a deep-learning method that predicts inter-protein contact maps and interface residues for an IDR and its partner, given their sequences. It uses sequence embeddings from the ProtT5 protein language model. Disobind outperforms state-of-the-art interface predictors for IDRs. It also outperforms AlphaFold-multimer and AlphaFold3 at multiple confidence cutoffs. Combining Disobind and AlphaFold-multimer predictions further improves the performance. In contrast to current methods, Disobind considers the context of the binding partner and does not depend on structures and multiple sequence alignments. Its predictions can be used to localize IDRs in large assemblies and characterize IDR-mediated interactions.

A record of this paper’s transparent peer review process is included in the supplemental information.

## Introduction

Intrinsically disordered proteins or regions (IDPs or IDRs) lack a well-defined three-dimensional structure in their monomeric state ^1,2^. They exist as an ensemble of interconverting conformers in equilibrium and hence are structurally heterogeneous^1,3,4^. Here, we use the term IDR for both IDPs and IDRs. The heterogeneity of IDRs provides several functional advantages including the ability to overcome steric restrictions, have a large capture radius, undergo functional misfolding, and interact with multiple partners ^5^. They play critical roles in cellular processes including signaling, intracellular transport, protein folding, and condensate formation ^4,5^.

IDRs are known to mediate a large number of protein-protein interactions ^6^. They exhibit considerable diversity in their binding modes within complexes ^7,8^. Upon binding to a partner protein, they may adopt an ordered state (disorder-to-order or DOR) or may remain disordered (disorder-to-disorder or DDR) ^7,9–11^. Moreover, depending on the binding partner, the same IDR may undergo DOR and DDR transitions ^10,11^. For example, IDRs with short linear motifs (SLiMs) may form multivalent interactions with their partner, exchanging between conformations where one or more motifs bind at a time ^11^. This heterogeneity in binding makes their structural characterization in complexes challenging.

Here, we develop a deep learning method to determine the binding sites of an IDR and an interacting partner protein given their sequences. Several methods have been developed for predicting interface residues, inter-protein contact maps, and structures of complexes formed by two proteins. Some of them rely on the availability of structures of the input proteins ^12–17^. Sequence-based methods that require a multiple sequence alignment (MSA) can be constrained by the low sequence conservation of IDRs ^18–21^. Other sequence-based methods for IDRs use physicochemical features or embeddings from protein language models (pLMs) but do not account for the partner protein ^22–25^. Yet, the identity of the partner protein is an important determinant of IDR interactions, as these are context-dependent. To this end, FINCHES predicts intermolecular interactions for two IDRs based on their chemical specificity, given their sequences. However, it is currently applicable to IDRs that remain disordered upon binding ^26^.

State-of-the-art methods like AlphaFold2 (AF2) and AlphaFold3 (AF3) can provide accurate predictions for a range of macromolecules ^18,20,27^. However, they tend to provide low-confidence predictions for IDRs. Even though the low-confidence regions predicted by AF2 often correspond to the presence of IDRs, the predicted structure cannot be considered a representative IDR structure ^28,29^. Recent studies showed that AlphaFold-multimer (referred to as AF2 hereon) could predict the structures of complexes where the IDR undergoes a DOR transition, including protein-peptide complexes and domain-motif complexes ^19,21,30^. However, the results may be sensitive to the inputs, such as the sequence fragment size, the fragment delimitation, and the alignment mode for the MSA ^19,21^. In general, predicting the binding interfaces for IDRs at high resolution remains a challenge for current methods ^31^.

Our method, Disobind, predicts the inter-protein contact maps and interface residues for an IDR and its partner, which could be an IDR or an ordered protein, from their sequences. Disobind is trained on experimental structures of IDRs in complexes. However, predicting inter-protein contact maps for IDRs presents several challenges. The structural data for IDRs in complexes is limited, the IDRs may retain their heterogeneity in complexes, and the inter-protein contact maps can be sparse. Here, we leverage the pLM ProtT5 to train our model with limited data, without the need for large, resource-intensive architectures. To address the problems associated with sparsity and heterogeneity, we predict coarse-grained inter-protein contact maps and interface residues. Coarse-graining reduces the resolution of the contact maps, making the prediction task easier. Disobind performs better than state-of-the-art interface predictors for IDRs and better than AF2 and AF3 across multiple confidence cutoffs used in this study. Combining the Disobind and AF2 predictions further improves the performance. We demonstrate the performance of Disobind+AF2 on several IDR complexes, including DOR and DDR complexes. Predictions from the method can be used to localize IDRs in integrative structures of large assemblies. They can be used to characterize protein-protein interactions involving IDRs, identify novel motifs, and modulate IDR-mediated interactions.

## Results

### Disobind dataset creation

Given a pair of protein sequences, at least one of which is an IDR, Disobind predicts binary inter-protein contact maps and interface residues for the pair (Fig 1). We compile our dataset by gathering experimental structures of IDR-containing complexes from an array of existing IDR databases and datasets, including DIBS, MFIB, FuzDB, PDBtot, PDBcdr, DisProt, IDEAL, and MobiDB, covering a range of IDR binding modes (Fig 1) (See Methods) ^9,32–36^. We first process the chains in the PDB structures of the IDR-containing complexes (Fig. 1). Missing residues in the structure cannot be used to infer contact maps reliably. Therefore, segments corresponding to missing residues are removed, resulting in fragments of the chain. Long chains are also fragmented to 100 residues for memory efficiency. Notably, this does not impose any sequence length restriction for Disobind; it merely determines the crop sizes used in training. At inference, no length restriction exists, and Disobind can be used for sequences of arbitrary lengths.

**Figure 1:**
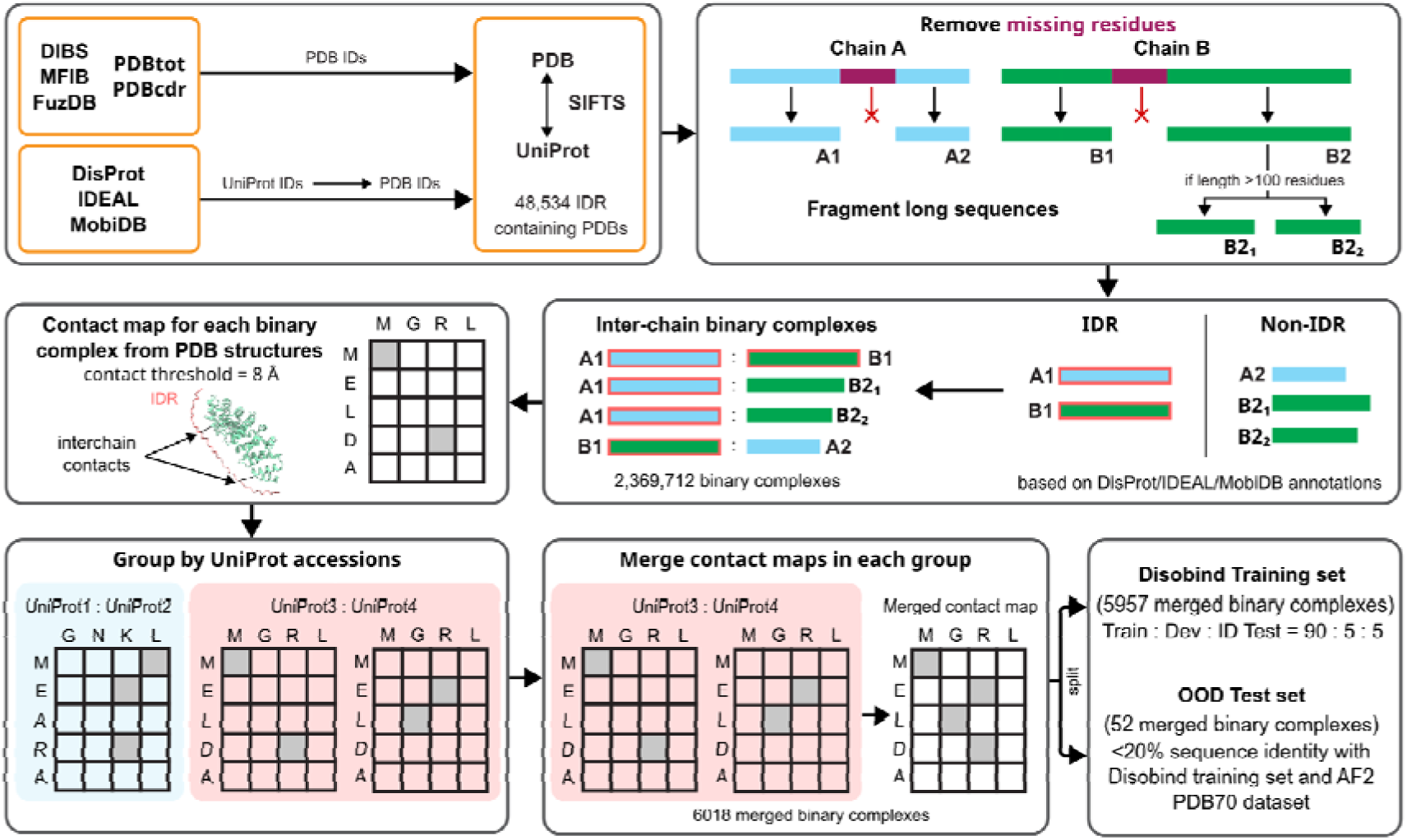
Disobind dataset creation pipeline. We compiled our dataset by gathering structures of IDR-containing complexes from DIBS, MFIB, FuzDB, PDBtot, PDBcdr, DisProt, IDEAL, and MobiDB. For each PDB chain, missing residues in the structures are excluded, resulting in sequence fragments. Further, we restrict the maximum length of a sequence fragment to 100 residues. We use annotations from DisProt, IDEAL, and MobiDB to classify a sequence fragment as an IDR and non-IDR. Next, we obtain inter-chain binary complexes comprising at least one IDR fragment (IDR with an ordered protein or both IDRs). Contact maps are obtained for all binary complexes which are subsequently grouped based on the UniProt accessions of the sequence fragment pair. The contact maps of all binary complexes in a group are merged, creating merged contact maps. A sequence fragment pair and its corresponding merged contact map comprise a merged binary complex. The merged binary complexes are redundancy reduced at a 20% sequence identity cutoff to create an out-of-distribution (OOD) test set. The remaining complexes comprise the Disobind training set which is split into train, dev, and in-distribution (ID) test sets.

We then designate the sequence fragments derived from the chains in the PDB structures as IDRs and non-IDRs. We identify disordered residues based on annotations supported by experimental evidence and sequence-based homology obtained from DisProt, IDEAL, and MobiDB. Based on these annotations, sequence fragments with more than 20% disordered residues are considered as IDR fragments, whereas the others are considered as non-IDR fragments (See Methods). The 20% threshold is used solely as a criterion for inclusion in our dataset and is not intended as a general definition of an IDR. This cutoff was chosen to ensure a sufficiently large dataset for training, while limiting the over-representation of ordered residues.

Next, we obtain inter-chain binary complexes from each PDB structure. A binary complex comprises sequence fragments from different chains, of which at least one is an IDR fragment (Fig. 1). Each such binary complex is associated with a contact map; we filter out complexes that do not contain any contacts. The binary complexes are then grouped based on the UniProt accessions of the corresponding sequence fragment pair.

We then merge all contact maps with overlapping sequences in a group using a logical OR operation to obtain merged contact maps (Fig. 1). A sequence fragment pair, along with its corresponding merged contact map, forms a merged binary complex (See Methods) (Fig. 1). The latter represents contacts formed across all available structures of complexes for a given sequence pair, accounting for the multiple conformations of an IDR in a complex. This simplified output representation avoids making any assumptions about binding mode, allowing it to be applicable to DOR and DDR cases. Further, this provides for a less sparse output representation compared to an ensemble of contact maps, while also mitigating the bias caused by having an unequal number of structures for different sequence fragment pairs. Disobind is trained on a dataset of merged binary complexes with the sequence fragments as input and the merged contact map as output.

We then create an out-of-distribution (OOD) test set comprising merged binary complexes where the input sequence fragments share less than 20% identity with the Disobind dataset and with the AlphaFold2 PDB70 dataset. The remaining complexes are further split into a train, validation, and an in-distribution (ID) test set. The ID and OOD test sets comprise 297 and 52 merged binary complexes respectively. Adding more complexes to the OOD test set may be achieved by increasing the sequence identity cutoff but would also increase similarity to the AF2 training dataset, which we attempted to avoid.

### Model architecture

Disobind uses a shallow feedforward neural network architecture with the sequence embeddings obtained from a protein language model as input (Fig 2). The input embeddings are projected to a lower dimension using a projection block. Next, in the interaction block, the projected embeddings are used to compute the outer product and outer difference, which are subsequently concatenated along the feature dimension. This allows the model to capture complementarity between the input sequences and yields an interaction tensor. This interaction tensor is processed by a multi-layer perceptron (MLP) for contact map prediction. For interface residue prediction, the interaction tensor is first reduced by computing row-wise and column-wise averages and further processed by an MLP. Finally, an output block provides element-wise sigmoid scores, with a score greater than 0.5 representing a contact or interface residue. The outputs are binary contact maps or binary interface residue predictions on the fragments. The same architecture is used to train separate models for the contact map and interface residue prediction tasks. The model is trained with the singularity-enhanced loss (SE loss) ^37^ using the AdamW optimizer with weight decay (Fig S1). This loss is a modified form of the binary cross-entropy (BCE) loss used for class-imbalanced training. We varied the hyperparameters of the network including the projection dimension, the number of layers in the MLP, and the SE loss parameters (Table S1-S4, Fig S1). We use recall, precision, and F1-score to evaluate model performance (See Methods).

**Figure 2:**
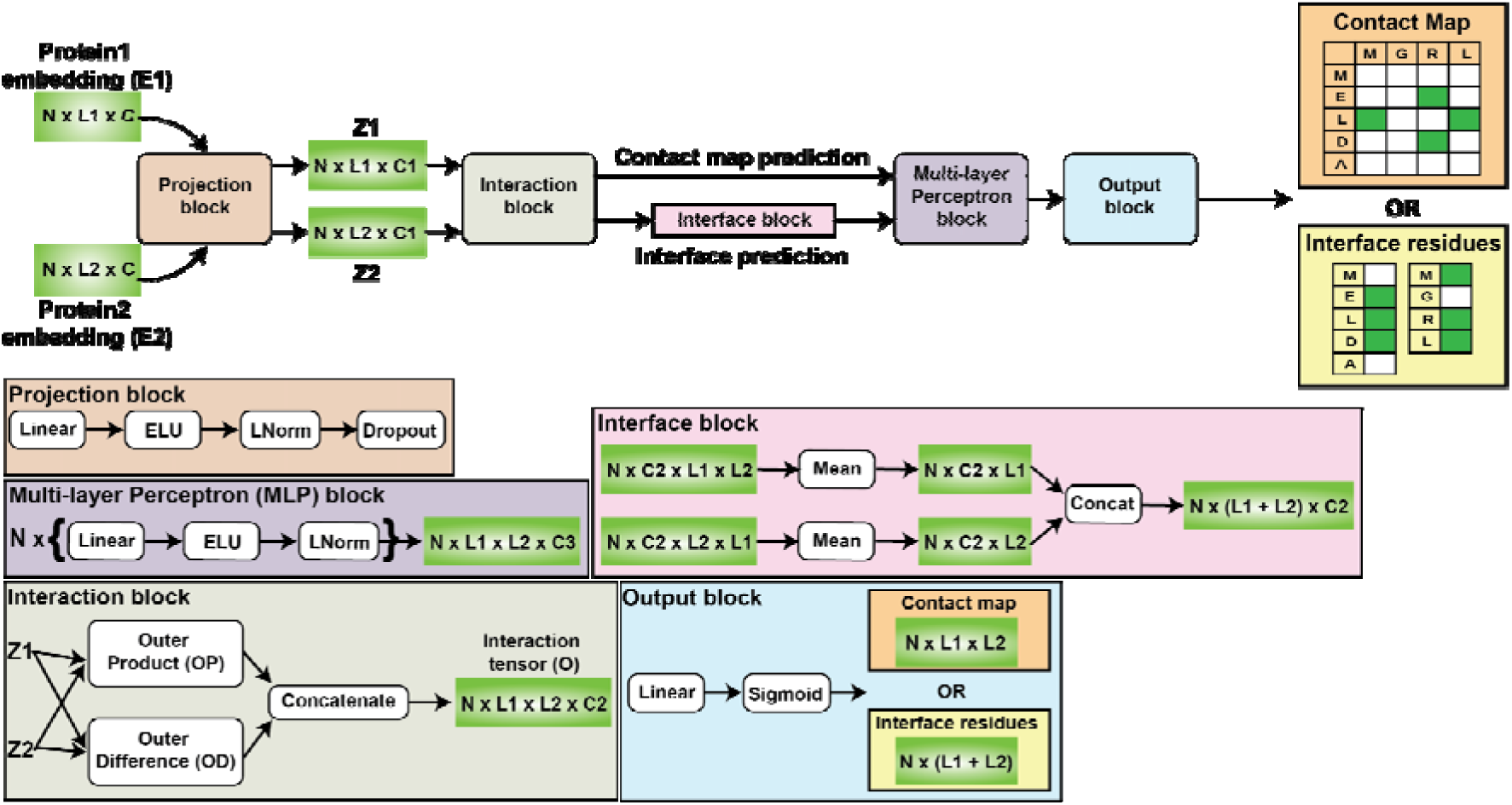
Disobind architecture. The inputs to Disobind are the sequence embeddings for the sequence pair obtained using a pLM. The sequence embeddings are projected to a lower dimension using a projection block. Next, in the interaction block, the projected embeddings are used to compute the outer product and outer difference, which is subsequently concatenated along the feature dimension, resulting in an interaction tensor. For contact map prediction, this interaction tensor is processed by a multi-layer perceptron (MLP). For interface residue prediction, the interaction tensor is reduced by computing row-wise and column-wise averages and further processed by an MLP. Finally, an output block provides element-wise sigmoid scores, with a score greater than 0.5 representing a contact or interface residue. Here, N, L, C represent the batch size, sequence length, and feature dimension respectively.

### Evaluating Disobind

#### Inter-protein contact map prediction

We first evaluate Disobind on predicting residue-wise inter-protein contact maps on the ID and OOD test sets. Disobind achieves an F1-score of 0.57 and 0.33 on the ID and OOD test sets respectively (Table 1 and Table S5, contact map prediction at coarse-grained (CG) resolution 1). It performs better than a random baseline (see Methods) on the OOD test set (Table 1). However, predicting residue-wise interprotein contact maps is challenging due to their sparsity (Table S6). This leads to a class imbalance, biasing the model to predict just the majority class, *i.e.,* 0’s or non-contacts. The heterogeneity of IDRs in complexes poses another challenge. For example, the multivalency of IDRs may result in different sets of contacts in different conformations with the same partner ^38^.

**Table 1:** Comparing Disobind, AF2, and AF3 performance. We compare the performances of Disobind, AF2, and AF3 on the OOD set for contact map and interface residue prediction for different coarse-grained (CG) resolutions. We also include a random baseline (RB) for comparison. For AF2 and AF3 we only consider the confident predictions (pLDDT >= 70 and PAE <= 5) at multiple ipTM cutoffs (0, 0.4, 0.75) for evaluation. The number after the ‘_’ represents the ipTM cutoff used.

| Model | Recall | Precision | F1 score | Recall | Precision | F1 score |
| --- | --- | --- | --- | --- | --- | --- |
|  | Contact map |  |  | Interface |  |  |
|  | Coarse Graining 1 |  |  |  |  |  |
| Disobind | 0.22 | 0.63 | 0.33 | 0.5 | 0.46 | 0.48 |
| RB | 0.51 | 0.01 | 0.02 | 0.5 | 0.26 | 0.34 |
| AF2_0.75 | 0.12 | 0.58 | 0.2 | 0.18 | 0.76 | 0.29 |
| AF3_0.75 | 0.02 | 0.41 | 0.04 | 0.02 | 0.43 | 0.04 |
| AF2_0.4 | 0.2 | 0.45 | 0.28 | 0.35 | 0.7 | 0.47 |
| AF3_0.4 | 0.08 | 0.33 | 0.13 | 0.18 | 0.6 | 0.27 |
| AF2_0.0 | 0.2 | 0.45 | 0.28 | 0.35 | 0.7 | 0.47 |
| AF3_0.0 | 0.08 | 0.31 | 0.13 | 0.18 | 0.57 | 0.27 |
|  | Coarse Graining 5 |  |  |  |  |  |
| Disobind | 0.38 | 0.61 | 0.47 | 0.73 | 0.56 | 0.63 |
| RB | 0.52 | 0.08 | 0.13 | 0.53 | 0.45 | 0.49 |
| AF2_0.75 | 0.15 | 0.58 | 0.24 | 0.2 | 0.79 | 0.32 |
| AF3_0.75 | 0.02 | 0.34 | 0.03 | 0.02 | 0.5 | 0.04 |
| AF2_0.4 | 0.29 | 0.52 | 0.37 | 0.43 | 0.76 | 0.54 |
| AF3_0.4 | 0.14 | 0.47 | 0.22 | 0.22 | 0.72 | 0.34 |
| AF2_0.0 | 0.29 | 0.52 | 0.37 | 0.43 | 0.76 | 0.54 |
| **AF3_0.0** | 0.14 | 0.45 | 0.22 | 0.23 | 0.7 | 0.34 |
|  | **Coarse Graining 10** |  |  |  |  |  |
| **Disobind** | 0.42 | 0.49 | 0.45 | 0.8 | 0.61 | 0.69 |
| **RB** | 0.54 | 0.17 | 0.26 | 0.48 | 0.52 | 0.5 |
| **AF2_0.75** | 0.16 | 0.62 | 0.25 | 0.2 | 0.83 | 0.32 |
| **AF3_0.75** | 0.02 | 0.4 | 0.04 | 0.03 | 0.59 | 0.05 |
| **AF2_0.4** | 0.32 | 0.57 | 0.41 | 0.44 | 0.8 | 0.57 |
| **AF3_0.4** | 0.16 | 0.52 | 0.25 | 0.24 | 0.78 | 0.36 |
| **AF2_0.0** | 0.32 | 0.57 | 0.41 | 0.44 | 0.8 | 0.57 |
| **AF3_0.0** | 0.16 | 0.51 | 0.25 | 0.24 | 0.75 | 0.37 |

#### Interface residue prediction

As an alternative to predicting inter-protein residue-wise contact maps, next, we predict interface residues for the IDR and its partner (Fig 3a). For two input sequence fragments of length *L_1_* and *L_2_*, this reduces the number of predicted elements from *L_1 ×_ L_2_* in the contact map prediction to *L_1_* + *L_2_* in the interface residue prediction (Fig 3b). This helps mitigate the class imbalance as the interface residue predictions are less sparse (Fig 3a, Fig 3b Table S6, interface residue prediction at coarse-grained (CG) resolution 1). Interface residue prediction is also less affected by the heterogeneity of IDRs in complexes. The prediction for each residue is simplified to determining whether it binds to the partner, rather than identifying the specific residues within the partner it interacts with.

**Figure 3:**
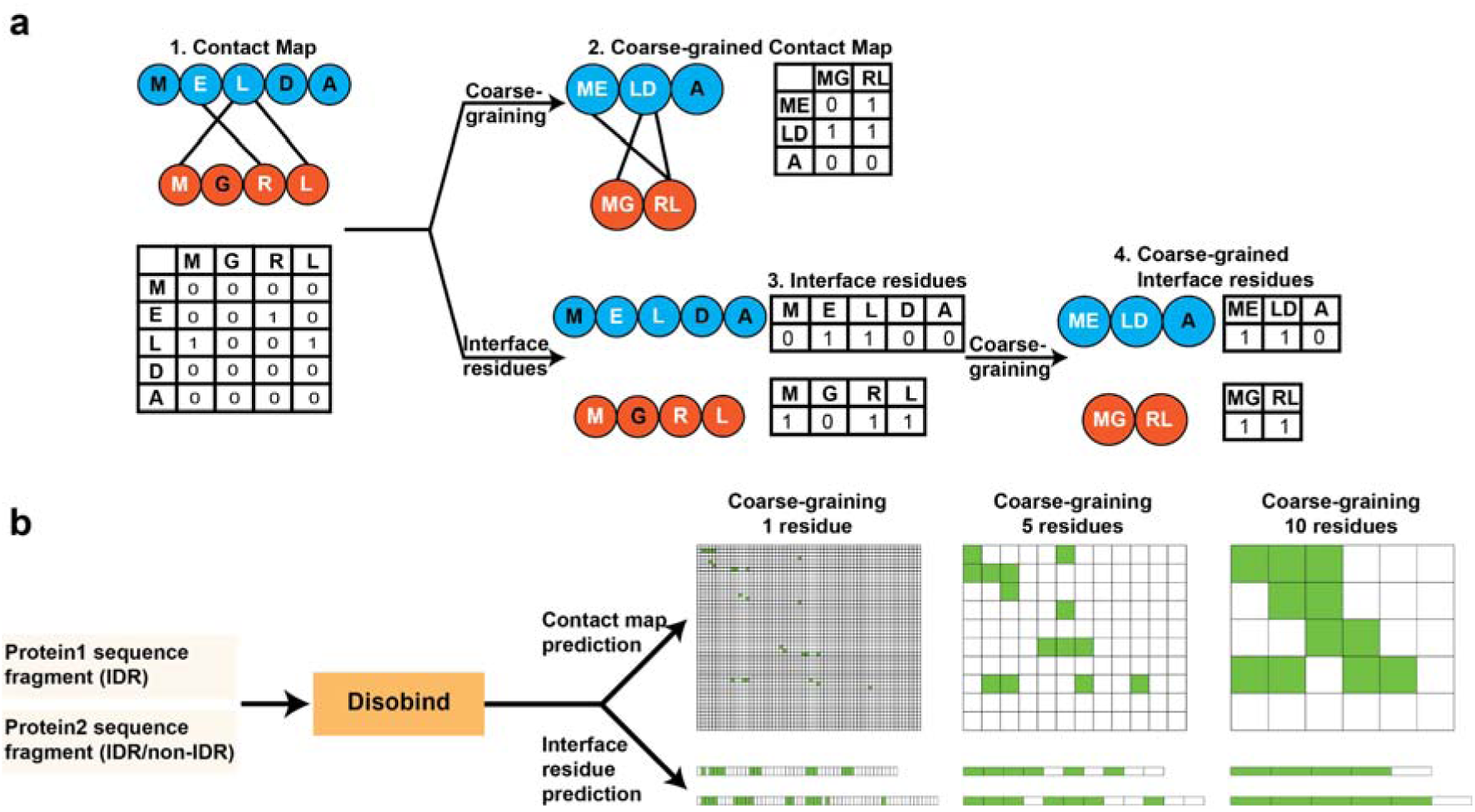
Predicting coarse-grained contact maps and interface residues. **(a)** We use coarse-graining and predict interface residues as an alternative to predicting residue-wise inter-protein contact maps. This addresses the problems associated with the multivalency of IDRs and the sparsity of inter-protein contact maps. **(b)** Disobind can predict inter-protein contact maps and interface residues at multiple coarse-grained resolutions (1, 5, 10).

As expected, Disobind performs better in interface residue prediction than in contact map prediction. It achieves an F1 score of 0.70 and 0.48 on the ID and OOD test sets respectively (Fig S2, Table 1 and Table S5, interface residue prediction at coarse-grained (CG) resolution 1). It performs better than a random baseline evaluated on the OOD test set (Table 1).

#### Coarse-graining improves the performance

To further improve the performance of our model, we predict coarse-grained contact maps and interface residues (Fig 3a, Fig 3b). Coarse-graining has been widely used for modeling biological systems ^39,40^. Coarse-graining over a set of residue pairs in a contact map or a set of residues in the interfaces further reduces the sparsity and class imbalance, and mitigates the problems posed by the multivalency of IDR interactions (Fig 3a, Fig 3b, Table S6). Notably, we coarse-grain over embeddings from pLMs. The latter provide context-aware, abstract representations of the sequence. Further, coarse-graining along the sequence is well-suited for the modular and patch-driven nature of IDR interactions, that are mediated by charge patterning, attractive sticker regions, conserved short linear motifs (SLiMs) and molecular recognition features (MoRFs), and/or less conserved low-complexity regions (LCRs)^7,41–43^.

We train Disobind to predict coarse-grained contact maps and coarse-grained interface residues at resolutions of 5 and 10 contiguous residues along the backbone. The coarse-grained embeddings are obtained by averaging the input embeddings ^44^. As expected, coarse-graining further improves the performance of Disobind on the ID and OOD test sets for both the contact map and interface residue prediction (Fig S2, Table 1, Table S5). For all cases, Disobind performs better than a random baseline on the OOD test set.

### Comparison to AlphaFold2 and AlphaFold3

We further compare the performance of Disobind with the state-of-the-art methods AF2 and AF3 on the OOD test set. For evaluating AF2 and AF3 outputs, we mask out interactions predicted at low confidence based on the interface-predicted template matching (ipTM) score, predicted local distance difference test (pLDDT), and predicted aligned error (PAE) metrics (see Methods).

#### Using different ipTM cutoffs for AF2 and AF3

The ipTM score is a metric provided by AF2 and AF3 to assess the confidence in the predicted interfaces in a complex ^18,27^. A prediction with an ipTM score greater than 0.75 is considered highly confident, while an ipTM lower than 0.6 indicates a likely failed prediction ^45,46^. We first evaluate the performance of AF2 and AF3 with an ipTM cutoff of 0.75. Disobind performs better than AF2 and AF3 for all tasks, *i.e.,* contact map and interface residue predictions across different coarse-grained resolutions (Table 1, ipTM = 0.75). Disordered residues, however, are known to impact the pTM and ipTM metrics negatively ^18,47^. Considering this, we relaxed the ipTM cutoff to 0.4 to assess the performance of AF2 and AF3, while keeping the per-residue metric, *i.e.* pLDDT and PAE cutoffs as is, guided by recent benchmarks and the AF3 guide ^18,30^. Here also, Disobind performs better than both AF2 and AF3 for both contact map and interface residue prediction (Table 1, ipTM = 0.4). AF3 performs worse than both Disobind and AF2. Furthermore, this trend remains more or less the same even when we do not apply an ipTM cutoff (Table 1, ipTM = 0.0).

Further, AF2 and AF3 are trained to output a distogram ^18,20,27^. The auxiliary distogram head is used to ensure that the learned internal pair representation accurately reflects the distribution of distances between the residues (See section Distogram head in AF2 and AF3 supplementary text). AF3 provides the contact probabilities derived from the distogram head in the output JSON file. We compared the performance of AF3 on the contact map and interface residue prediction task on the OOD test set using residue-level contact maps derived in two ways: 1) from the predicted AF3 structure, and 2) from the contact probabilities from the distogram head. The results from both methods are in good agreement, indicating that either of them can be used to obtain contact maps from AF3 predictions (Table S7).

#### AF2 performs better than AF3

Similar to Disobind, coarse-graining improves the performance of both AF2 and AF3 for both contact map and interface predictions (Table 1). Across all tasks, AF2 performs better than AF3. Moreover, AF3 predictions have lower confidence as measured by their ipTM scores (Fig S3). For 35 of 52 OOD test set entries, the ipTM score of the best AF2 model was greater than that of the best AF3 model. Overall, only 8 of 52 predictions from AF2 and 2 of 52 predictions from AF3 had a high confidence prediction, *i.e.*, ipTM score higher than 0.75.

#### Combining Disobind and AlphaFold2 predictions

As AF2 performed better than AF3, we asked if combining the predictions from Disobind and AF2 could further improve over either method. We combine the predictions from Disobind and AF2 using a logical OR operation and evaluate the performance on the OOD test set. The combined model, “Disobind+AF2”, performs better than either of the methods (Table 2, see also Supplementary Text).

**Table 2:** Evaluating the performance of Disobind+AF2 on the OOD test set. We combine Disobind and AF2 predictions using a logical OR operation and evaluate the performance on the OOD test set for contact map and interface residue prediction. For AF2 we consider the confident predictions (pLDDT >= 70 and PAE <= 5) for evaluation. No ipTM cutoff is applied for AF2 and Disobind+AF2 predictions.

| Model | Recall | Precision | F1 score | Recall | Precision | F1 score |
| --- | --- | --- | --- | --- | --- | --- |
|  | Contact map |  |  | Interface |  |  |
|  | Coarse Graining 1 |  |  |  |  |  |
| Disobind | 0.22 | 0.63 | 0.33 | 0.5 | 0.46 | 0.48 |
| AF2_0.0 | 0.2 | 0.45 | 0.28 | 0.35 | 0.7 | 0.47 |
| Disobind+AF2 | 0.33 | 0.49 | 0.4 | 0.6 | 0.46 | 0.52 |
|  | Coarse Graining 5 |  |  |  |  |  |
| Disobind | 0.38 | 0.61 | 0.47 | 0.73 | 0.56 | 0.63 |
| AF2_0.0 | 0.29 | 0.52 | 0.37 | 0.43 | 0.76 | 0.54 |
| Disobind+AF2 | 0.49 | 0.5 | 0.5 | 0.78 | 0.54 | 0.64 |
|  | Coarse Graining 10 |  |  |  |  |  |
| Disobind | 0.42 | 0.49 | 0.45 | 0.8 | 0.61 | 0.69 |
| AF2_0.0 | 0.32 | 0.57 | 0.41 | 0.44 | 0.8 | 0.57 |
| Disobind+AF2 | 0.55 | 0.46 | 0.5 | 0.86 | 0.61 | 0.71 |

### Performance by residue types

Next, we assess the performance of Disobind, AF2, and Disobind+AF2 by residue type to examine which regions each method tends to perform better on (See Methods). Disobind outperforms AF2 on both the contact map and interface residue prediction in disordered regions, whereas AF2 performs slightly better than Disobind for ordered regions (Table S8). Importantly, the combined model Disobind+AF2 performs better for both disordered and ordered regions, likely because the two models learn complementary information by focusing on different regions.

Further, Disobind predictions are more accurate than AF2 for disorder-promoting and hydrophobic, whereas AF2 and Disobind perform similarly for aromatic and polar residues (Table S9). Disobind is more accurate than AF2 in predicting contacts and interface residues involving LIPs (linear interacting peptides) (Table S9). The combined model is better across all residue types (Table S9).

Lastly, we assess the performance of Disobind, AF2, and Disobind+AF2 on the 17 OOD complexes containing long IDRs, *i.e.,* complexes where 100% of the residues in the IDR fragment are annotated as disordered (Table S10). Disobind and AF2 perform comparably at contact map prediction, while Disobind performs slightly better at interface prediction. Disobind+AF2 achieves the best results overall for both contact map and interface residue prediction (Table S10).

### Comparison with interface predictors for IDRs

We compared Disobind with partner-independent interface predictors for IDRs, including AIUPred, MORFchibi, and DeepDISOBind, which are comparable to the state-of-the-art in the Critical Assessments of protein Intrinsic Disorder prediction (CAID2) ^25,48–50^. For a fair comparison, we evaluated interface residue predictions for only the IDR protein in the OOD set (See Methods). Disobind outperforms AIUPred, MORFchibi, and DeepDISOBind at predicting interface residues for the IDR across all coarse-grained resolutions (Table 3).

**Table 3:** Comparison with interface predictors for IDRs. We compare the performance of Disobind with interface predictors for IDRs, including AIUPred, MORFchibi, and DeepDISOBind, on the IDR sequences from the OOD test set.

| Model | Recall | Precision | F1 score |
| --- | --- | --- | --- |
|  | **Interface residue prediction, CG1** |  |  |
| **Disobind** | 0.5 | 0.49 | 0.5 |
| **Aiupred** | 0.36 | 0.38 | 0.37 |
| **Morfchibi** | 0.97 | 0.24 | 0.39 |
| **Deepdisobind** | 0.22 | 0.45 | 0.29 |
|  | **Interface residue prediction, CG5** |  |  |
| **Disobind** | 0.73 | 0.56 | 0.63 |
| **Aiupred** | 0.43 | 0.58 | 0.49 |
| **Morfchibi** | 0.96 | 0.41 | 0.58 |
| **Deepdisobind** | 0.21 | 0.73 | 0.33 |
|  | **Interface residue prediction, CG10** |  |  |
| **Disobind** | 0.8 | 0.61 | 0.69 |
| **Aiupred** | 0.43 | 0.61 | 0.51 |
| **Morfchibi** | 0.95 | 0.5 | 0.66 |
| **Deepdisobind** | 0.2 | 0.79 | 0.31 |

### Evaluating Disobind for protein-protein interaction prediction

We evaluated Disobind for predicting protein-protein interactions (PPI), *i.e.* to predict if two input protein sequences will bind or not. We used the five PPI test datasets for IDRs provided by ^51^. The dataset comprises UniProt accessions for interacting and non-interacting protein pairs. This includes a total of 9993 protein pairs. The dataset was pre-processed by removing protein pairs for which the UniProt accessions were part of the Disobind training set, the UniProt sequence could not be downloaded, and the sequence length was greater than 10000 residues. This resulted in a total of 2355 protein pairs. Disobind interface residue predictions at coarse-grained resolution 1 were obtained for all the protein pairs. To obtain the PPI predictions from interface residue predictions, we aggregate the residue-level predictions using two ways: (i) maximum sigmoid score across both proteins, and (ii) average sigmoid score across both proteins, and evaluate the performance using the Area Under the Receiver Operator Curve (AUROC). Disobind achieves an AUROC of 0.48 and 0.5, for the two ways, respectively, indicating a random prediction.

### Input Embeddings

Next, we investigate the effect of using embeddings from various pLMs and the different embedding types on the performance of Disobind.

#### Protein language models allow for a shallow architecture

Protein language models (pLMs) provide context-aware representations of protein sequences that facilitate training models for downstream tasks like contact map prediction *via* transfer learning. They allow using a shallow downstream architecture when training with limited data ^22,52^. Several pLMs have been developed including the ESM series of models ^53^, ProstT5 ^54^, ProtT5 ^44^, ProtBERT ^44^, and ProSE ^55^.

The pLMs differ in the model architecture, the training datasets, and the training regime. These differences can affect their ability to learn contextual representations for protein sequences. ProSE uses a bidirectional LSTM trained on UniRef protein sequences and structures with a multitask loss combining similarity and contact prediction objectives ^55^. The other pLMs use variants of the transformer architecture. ProtBERT uses the Bidirectional Encoder Representations from Transformers (BERT) architecture, an encoder-only model, trained on the UniRef100 and BFD100 datasets. It is trained to reconstruct corrupted tokens given the context of the neighbouring tokens ^44^. ProtT5 uses the Text-to-Text Transfer Transformer (T5) architecture, an encoder-decoder model, trained on the UniRef50 and BFD100 datasets. In contrast to the BERT-style training objective, ProtT5 is trained to reconstruct spans of corrupted tokens ^44,56^. ProstT5 is a newer version of ProtT5, fine-tuned with AlphaFold2 predicted protein structures ^57^.

Disobind uses embeddings from ProtT5, one of the best performing pLMs in our comparison (Table S11). The T5-based models, ProtT5 and ProstT5, perform similarly, but outperform the BERT-styled ProtBERT and LSTM-based ProSE models. Due to limitations in memory, we could not generate ESM2-650M-global embeddings for the entire training set. Hence, we do not include ESM in our comparison.

#### Global embeddings perform better than local embeddings

We then evaluate whether models trained with global sequence embeddings perform better than those with local embeddings. Local embeddings are obtained by providing the sequence of the fragment as input to the pLM. Global embeddings, in contrast, are obtained by providing the entire protein sequence corresponding to the fragment as input to the pLM, followed by extracting the embeddings corresponding to the fragment (Fig S4). Models with global embeddings perform better than those with local embeddings across all the pLMs we tested (Table S11). Using the global embeddings provides the context of the complete protein sequence corresponding to the fragment, plausibly resulting in better performance. This context may contain information relevant to binding: for example, the binding of IDRs is known to be affected by the flanking regions ^8^.

### Case studies

IDRs enable dynamic interactions, molecular recognition, and functional adaptability, making them a crucial component of various signaling and regulatory pathways. We used Disobind+AF2 for predicting interface residues for several IDR complexes, as it performs better than the other models (Fig 4, Fig S5). Here, we discuss in detail two cases in which the IDR becomes ordered upon binding (DOR) and two cases where the IDR remains disordered (DDR). A larger set of NMR examples of mostly DDR cases from MobiDB, PED, BMRB, and LLPSdb are also shown separately (Fig S5)^35,58–60^.

**Figure 4:**
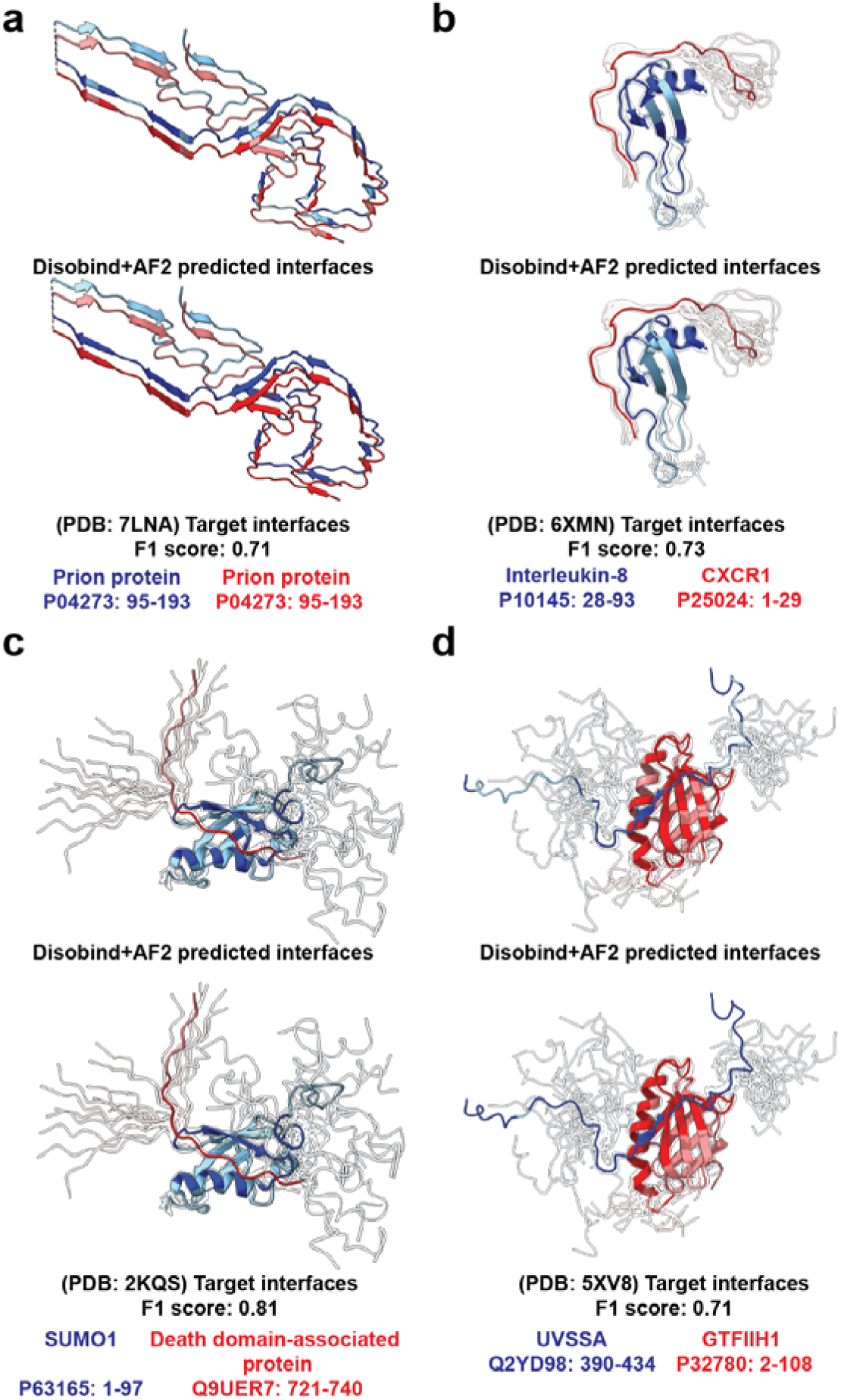
Case studies visualizing Disobind+AF2 predictions. The figures visualize the Disobind+AF2 predicted and target interface residues for both proteins in complexes where IDRs undergo a DOR (panels a and b) or DDR (panels c and d) transition. The protein chains are coloured in blue and red, where the lighter shade represents the chain and the darker shade represents the interface residues for the respective chains. For all NMR structures, the ensemble is represented with reduced transparency in the background, whereas the first model is shown in the foreground. The PDB IDs, protein names, and corresponding UniProt accession, along with the residue numbers for which the predictions were made, are shown for all complexes.

The cellular prion protein (PrP^C^) plays an important role in several cellular processes, including synapse formation, regulating circadian rhythm, and maintaining ion homeostasis (Kraus et al., 2021; Sigurdson et al., 2019). It is known to convert to an infectious PrP^Sc^ form, which forms amyloid fibrils associated with several diseases, including Creutzfeldt-Jakob disease and bovine spongiform encephalopathy (Kraus et al., 2021; Sigurdson et al., 2019). PrP^C^ has an N-terminal disordered region and a primarily α-helical C-terminal region, whereas the conversion to PrP^Sc^ involves significant structural changes, resulting in a β-sheet-rich structure. Disobind+AF2 could predict the interface residues in the β-sheet-rich PrP^Sc^ dimer with an F1-score of 0.71 (Fig 4a).

Chemokines are small secretary molecules that play several important roles, including immune cell trafficking, wound healing, and lymphoid cell development (Rossi & Zlotnik, 2000; Sepuru et al., 2020). Dysregulation of chemokine release is associated with several diseases, including chronic pancreatitis, inflammatory bowel disease, and psoriasis (Rossi & Zlotnik, 2000; Sepuru et al., 2020). Chemokines bind to chemokine receptors that belong to the G-protein coupled receptor (GPCR) family. Tha latter contains an N-terminal disordered region that becomes structured and adopts an extended conformation upon binding. Disobind + AF2 reliably predicts the interface residues for the chemokine Interleukin-8 binding to the N-terminus of its cognate receptor, CXCR1 with an F1 score of 0.73 (Fig 4b).

The death domain-associated protein 6 (Daxx) is a transcriptional coregulator that plays a key role in apoptosis ^61,62^. It contains a C-terminal disordered region that attains an extended conformation upon binding to its partner SUMO-1 ^61^. This interaction is crucial for its localization to nuclear PML bodies, the absence of which leads to disruption of PML nuclear bodies as seen in acute myelocytic leukaemia ^62^. Disobind+AF2 could reliably capture interface residues in this region, achieving an F1-score of 0.81 (Fig 4c).

UV-stimulated scaffold protein A (UVSSA) is a crucial component for transcription-coupled nucleotide excision repair (TC-NER) pathway ^63,64^. Disruption of the TC-NER pathway is associated with various diseases, including xeroderma pigmentosa and UV-sensitive syndrome ^63^. In complex with USP7, UVSSA is involved in processing the stalled RNA pol II and recruiting TFIIH. A short acidic region in UVSSA is an IDR and interacts with the N-terminal PH domain in the p62 subunit of the TFIIH complex, which is crucial for the recruitment of TFIIH to the TCR site ^63^. Disobind+AF2 was able to capture the interface residues in this case with an F1score of 0.71 (Fig 4d). In summary, across all the cases, our combined model achieves high accuracy in predicting interface residues for IDR complexes.

## Discussion

Here, we discuss about the uses and limitations of Disobind, challenges involved, the performance of AlphaFold, and future directions. Our combined model, Disobind+AF2, can be readily extended to any current or future structure prediction method that provides confidence metrics such as pLDDT and PAE for the predicted structures. Predictions from Disobind+AF2 may be used to improve the localization of IDRs in integrative models of large assemblies, our primary motivation for developing the method. Macromolecular assemblies contain significant portions of IDRs, for example, the Fg Nups in the nuclear pore complex, the MBD3-IDR in the nucleosome remodeling and deacetylase (NuRD) complex, and the N-terminus of Plakophilin1 in the desmosome ^65–67^. These regions typically lack data for structural modeling, resulting in integrative models of poor precision ^66,67^. The predictions from Disobind+AF2 can be used in integrative modeling methods such as IMP, HADDOCK, and Assembline as inter-protein distance restraints ^68–70^. Further, the predicted contacts can be combined with molecular dynamics (MD) simulations to provide mechanistic insights into their dynamic behaviour. Additionally, our method can be used to characterize IDR-mediated interactions across proteomes, helping identify new binding motifs linked to subcellular localization or function. It may also guide modulation of IDR interactions, for instance, by suggesting plausible mutations.

However, our method also has several limitations. First, it is limited to binary IDR-partner complexes. For complexes formed by three or more subunits, binary predictions from Disobind would need to be combined. Second, it assumes that the IDR and its partner are known to bind. Disobind cannot reliably distinguish binders from non-binders. Third, the accuracy of the predictions depends on the ability of the pLM to provide accurate representations of the IDR and its partner. Finally, our method cannot be used to assess the effects of post-translational modifications as the pLM used does not distinguish post-translationally modified amino acids.

The aforementioned limitations in part arise due to the challenges in predicting contact maps and interface residues for IDRs in complex with a partner. First, a limited number of experimental structures are available for IDRs in complexes ^22,38^. Second, IDRs adopt an ensemble of conformations, and the available structures may only partially capture this conformational diversity. Third, inter-protein contact maps are typically sparse, with only a few residues forming contacts. Although not sufficient, we gather all available structures for IDRs in complexes. With these structures, we create a dataset of merged binary complexes for training Disobind. Using merged binary complexes helps overcome the issue of sparsity associated with training on an ensemble of contact maps. Predicting interface residues instead of contacts and using coarse-graining helps overcome the challenges associated with the multivalency of IDRs and the sparsity of inter-protein contact maps (Fig 3a).

Several studies indicate that the low-confidence regions in AF2 predictions overlap with the presence of disordered regions, although these low-confidence regions cannot be considered as a representative conformation for IDRs ^28,29,71^. Despite the success of AF2 and AF3 for ordered proteins, their performance on IDR complexes remains limited, as shown in benchmarks ^19,28,30,72^. This could potentially be due to several reasons. First, IDRs show low sequence conservation, resulting in poor quality MSAs ^7^. Second, IDRs are best described by an ensemble of conformations which capture their dynamic behaviour. AF2 and AF3, however, predict a single best structure, which may not be representative of IDRs ^29^. Further, the diversity of binding modes of IDRs in complexes, *e.g.,* DOR and DDR, poses a significant challenge. In particular, AlphaFold predictions are less accurate for fuzzy complexes or DDR complexes, compared to DOR complexes ^28,30^. Third, the confidence metrics of AF2 and AF3 may not reliably reflect structural accuracy ^73,74^. Particularly, the presence of IDRs is known to negatively impact the confidence metrics (Dunbrack, 2025; Magana & Kovalevskiy, 2024; Omidi et al., 2024; Varga et al., 2024).

On our OOD test set, most of the predictions from AF2 and AF3 had low confidence, with an ipTM score lower than 0.75. Notably, compared to AF2, AF3 predictions were less accurate and had lower confidence. It is possible that the confidence cutoffs used for ordered proteins do not apply to IDRs and predictions involving the latter might require different cutoffs. A recent benchmark suggested a lower ipTM score of 0.4 for assessing interfaces involving IDRs ^30^. Further, the per-residue metrics such as pLDDT and PAE may be more relevant for assessing AF2 and AF3 predictions than the global metrics such as the ipTM and pTM, as is also suggested by a recent benchmark and the AF3 guide ^18,30^. Thus, AlphaFold confidence metrics might need to be interpreted differently for IDRs, and this requires rigorous benchmarking ^76–78^.

AF2 has been shown to successfully predict the structures of protein-peptide complexes and domain-motif complexes, although the results could be sensitive to the sequence fragment size, fragment delimitation, and the MSA alignment mode^19,21^. Some studies show that MSA subsampling^79^, clustering the sequences in the MSA ^80^, and using sliding fragments as input to AF2 ^19^ result in better predictions and can be used to predict multiple conformations. In our comparison, we did not explore these strategies, though they may improve the model predictions.

We conclude by highlighting potential future directions. One of the major roadblocks in training methods such as Disobind is the lack of data. More experimental structures of IDRs in complexes would be valuable ^22,38^. Whereas IDR ensembles derived from MD can be used, generating MD ensembles for IDR complexes is computationally expensive and challenging, and the existing databases like PED contain very few such ensembles for IDR complexes ^58,81^. Alternatively, deep generative models can be used to generate ensembles for IDRs in complexes ^81–83^. However, the current methods are limited to generating ensembles for monomers.

Compared to contact maps, distograms model the probability distribution across multiple distance ranges for each residue pair and thus provide a richer representation of structural heterogeneity for IDRs. However, predicting distograms for IDRs presents some non-trivial challenges. The limited structural data available for IDRs in complexes introduces strong biases toward a few observed conformations, making it difficult to learn representative distance distributions that capture the dynamic nature of IDRs. Furthermore, a distogram representation exacerbates the data sparsity problem due to the increased dimensionality of the predicted output from ℝ*^N×N^* for a contact map to ℝ*^N×N×d^* in a distogram, where *N* is the protein length and *d* is the number of distance bins. Thus, developing methods to predict distograms using sparse data for IDR complexes presents a promising avenue for future research.

Disobind and similar methods can be further enhanced by improving the existing pLMs to provide better representations for IDRs plausibly by incorporating physical priors and/or structural information ^81,84,85^. Additionally, these methods could be extended to predict protein-protein interactions (PPIs) involving IDRs, and aid in the design of IDR binders. The behaviour of IDRs within cells remains largely unexplored. Methods like Disobind are expected to facilitate an improved understanding of the interactions, function, and modulation of IDRs.

## Resource Availability

### Lead Contact

Further information and requests for resources and reagents should be directed to and will be fulfilled by the Lead Contact, Shruthi Viswanath.

### Materials Availability

This study did not generate new materials.

### Data and Code Availability

● Source data statement: The datasets created in this manuscript have been deposited to Zenodo at https://doi.org/10.5281/zenodo.14504762. The deposition contains all datasets (train, dev, ID, OOD) along with the input sequences and target contact maps, AF2 and AF3 predictions for the OOD test set, and the PDB structures, SIFTS mappings, PDB API files, and Uniprot Sequences used to create the datasets.
● Code statement: The Disobind model and scripts to create datasets, train the model, use Disobind/Disobind+AF2, and perform analysis are available at GitHub https://github.com/isblab/disobind.
● Any additional information required to reanalyze the data reported in this paper is available from the lead contacts upon request.

## Acknowledgement

We thank ISB Lab members Shreyas Arvindekar, Muskaan Jindal, Omkar Golatkar, and Mubashira KP for their useful comments on the manuscript. We also thank Aditi S. Pathak, Revathy Menon, Sarthak Joshi, Jagannath Mondal, Anand Srivastava, Vinothkumar KR, and Shachi Gosavi for valuable suggestions on the manuscript. We thank Omkar Golatkar for help in preparing Fig. 1. Molecular graphics images were produced using the UCSF Chimera and UCSF ChimeraX packages from the Resource for Biocomputing, Visualization, and Informatics at the University of California, San Francisco (supported by NIH P41 RR001081, NIH R01-GM129325, and National Institute of Allergy and Infectious Diseases).

## Author contribution

Conceptualization: K.M., S.V.

Reading and synthesis: K.M., S.V.

Data curation: K.M.

Methodology: K.M., V.U., S.V.

Writing – original draft: K.M.

Writing – review & editing: K.M., V.U., S.V.

Resources, Supervision, Funding acquisition: S.V.

## Funding

This work has been supported by the following grants: Department of Atomic Energy (DAE) TIFR grant RTI 4006, Department of Science and Technology (DST) SERB grant SPG/2020/000475, and Department of Biotechnology (DBT) grant BT/PR40323/BTIS/137/78/2023 from the Government of India to S.V.

## Declaration of interests

None declared.

## Diversity and inclusion statement

One or more of the authors in this paper self-identifies as an underrepresented ethnic minority in their field of research or within their geographical location. One or more of the authors in this paper self-identifies as a gender minority in their field of research.

## Declaration of generative AI and AI-assisted technologies

During the preparation of this work, the authors used ChatGPT to refine the wording. After using this tool/service, the authors reviewed and edited the content as needed and take full responsibility for the content of the publication.

## Supplemental information titles and legends

Supplementary material PDF – Figures S1-S5 and Tables S1-S11.

Table S1: Hyperparams.xlsx – Excel file containing hyperparameters for model training.

## STAR Methods

### Resource Availability

#### Materials Availability

This study did not generate new materials.

## Key resources table

| REAGENT or RESOURCE | SOURCE | IDENTIFIER |
| --- | --- | --- |
| **Antibodies** |  |  |
| NA |  |  |
| NA |  |  |
| NA |  |  |
| NA |  |  |
| **Bacterial and virus strains** |  |  |
| NA |  |  |
| NA |  |  |
| NA |  |  |
| NA |  |  |
| **Biological samples** |  |  |
| NA |  |  |
| NA |  |  |
| NA |  |  |
| NA |  |  |
| **Chemicals, peptides, and recombinant proteins** |  |  |
| NA |  |  |
| NA |  |  |
| NA |  |  |
| NA |  |  |
| **Critical commercial assays** |  |  |
| NA |  |  |
| NA |  |  |
| NA |  |  |
| NA |  |  |
| **Deposited data** |  |  |
| DIBS | Schad E. et. al. (2018) Bioinform. | https://dibs.enzim.ttk.mta.hu/ |
| MFIB | Fichó E. et. al. (2017) Bioinform. | https://mfib.pbrg.hu/ |
| FuzDB | Miskei M. et. al. (2017) NAR | https://fuzdb.org/ |
| PDBtot, PDBcdr | Miskei M. et. al. (2020) JMB | https://doi.org/10.1016/j.jmb.2020.02.017 |
| MobiDB | Piovesan D. et. al. (2024) NAR | https://mobidb.org/ |
| DisProt | Aspromonte M.C. et. al. (2024) NAR | https://disprot.org/ |
| IDEAL | Fukuchi S. et. al. (2014) NAR | https://www.ideal-db.org/ |
| PDB | Berman H.M. et. al. (2000) NAR | https://www.rcsb.org/ |
| UniProt | The UniProt Consortium (2025) NAR | https://www.uniprot.org/ |
| **Experimental models: Cell lines** |  |  |
| NA |  |  |
| NA |  |  |
| NA |  |  |
| NA |  |  |
| **Experimental models: Organisms/strains** |  |  |
| NA |  |  |
| NA |  |  |
| NA |  |  |
| NA |  |  |
| NA |  |  |
| **Oligonucleotides** |  |  |
| NA |  |  |
| NA |  |  |
| NA |  |  |
| NA |  |  |
| **Recombinant DNA** |  |  |
| NA |  |  |
| NA |  |  |
| NA |  |  |
| NA |  |  |
| **Software and algorithms** |  |  |
| SIFTS | Velanker S. et. al. (2013) NAR | 10.1093/nar/gks1258 |
| AlphaFold2 | Jumper J. et. al. (2021) Nature | https://github.com/google-deepmind/alphafold |
| AlphaFold3 server | Abramson J. et. al. (2024) Nature | https://alphafoldserver.com/ |
| MMSeqs2 | Steinegger M. et. al. (2017) Nat. Biotechnol. | https://github.com/soedinglab/MMseqs2.git |
| Pytorch v 2.0.1 |  |  |
| CUDA v 11.8 |  |  |
| AIUPred | Erdős G et. al. (2025) JMB | https://doi.org/10.1093/nar/gkae385 |
| MORFchibi 2.0 | Malhis N and Gsponer J (2025) bioRxiv | https://doi.org/10.1101/2025.01.31.635962 |
| DeepDISOBind | Zhang F et al (2021) Brief. in Bioinform. | https://doi.org/10.1093/bib/bbab521 |
| ProtT5, ProtBERT | Elnaggar A. et. al. (2023) IEEE PAMI | https://github.com/agemagician/ProtTrans.git |
| ProstT5 | Heinzinger M. et. al. (2024) NAR genom. bioinform. | https://github.com/mheinzinger/ProstT5.git |
| ProSE | Bepler and Berger (2019) ICLR | https://github.com/tbepler/prose.git |
| Disobind | This study | https://doi.org/10.5281/zenodo.14504762 |
| **Other** |  |  |
| Disobind GitHub | This study | https://github.com/isblab/disobind |
| Disobind Zenodo | This study | https://doi.org/10.5281/zenodo.14504762 |

## Experimental model and study participant details

Not applicable

## Method details

### Creating the dataset for Disobind

#### Gathering PDB structures of IDRs in complexes

First, we gathered the available PDB entries for protein complexes containing IDRs from databases on structures of disordered proteins: DIBS ^36^, MFIB ^33^, and FuzDB ^9^ (Fig 1). We also included PDB entries from the PDBtot and PDBcdr datasets ^10^. PDBtot consists of IDRs that either undergo a DOR or DDR transition, whereas PDBcdr consists of IDRs that undergo both DOR and DDR transitions depending upon the partner. We considered all PDB entries associated with each database. This resulted in a set of 4252 PDB entries. PED is another source of conformational ensembles of IDRs ^58^. However, PED entries were not used: several PED entries were already incorporated into our pipeline and the remaining entries correspond to monomers.

We then supplemented this set by querying the PDB for additional complexes containing IDRs. For this, we first obtained Uniprot identifiers (IDs) of proteins containing IDRs from DisProt ^32^, IDEAL ^34^, and MobiDB ^35^ (Fig 1). Specifically, from Disprot, we obtained all UniProt IDs corresponding to each DisProt entry. From IDEAL, we obtained UniProt IDs for entries annotated as “verified ProS”. These sequences are experimentally verified to be disordered in isolation and ordered upon binding. From MobiDB, we obtained UniProt IDs for entries annotated “curated-disorder-priority”, “homology-disorder-priority”, “curated-lip-priority”, and “homology-lip-priority”. These annotations correspond to sequences with curated experimental evidence for disorder (“curated-disorder”), their homologs (“homology-disorder”), sequences with experimental evidence for disorder with binding motifs such as SLIMs (“curated-lip”), and their homologs (“homology-lip”), respectively. Homology-based annotations were incorporated due to the lack of sufficient experimentally curated data available for IDRs. Importantly, the annotations were based on the source with the highest confidence (“priority”), and annotations based on indirect sources of evidence or predictions were not considered. Choosing the highest confidence annotation may reduce false positive “disorder” or “lip” annotations that may arise from homology-based evidence.

A total of 357734 unique Uniprot IDs across MobiDB, Disprot, and IDEAL were obtained. The PDB REST API was queried to obtain all the PDB entries associated with a given Uniprot ID, resulting in 48534 PDB entries (Table S1) ^86^. Combining these with the PDB entries obtained in the previous step, we obtained 50294 unique PDB entries. We note that MobiDB now provides the PDB IDs for IDR-containing complexes, however, this functionality was not available when this project started.

#### Defining binary complexes containing IDRs

The above PDB entries were downloaded using the Python requests library and mapped to the sequences in UniProt using the SIFTS (Structure Integration with Function, Taxonomy, and Sequence) tool ^87^. Entries with an obsolete PDB ID and those lacking a SIFTS mapping were removed, and deprecated PDB IDs were replaced with superseding PDB IDs. Further, entities corresponding to chimeric proteins or non-protein molecules, those having an obsolete UniProt ID, and those containing more than 10000 residues were removed.

Next, for each PDB entry, we obtained sequence fragments by removing missing residues from the sequence of each chain. The length of these fragments was kept to between 20 and 100 residues. Fragments longer than 100 residues were further divided due to constraints on GPU memory, that limited the scalability of the model architecture. We then identified the disordered residues in these fragments (Fig 1). This was achieved by cross-referencing the UniProt mapping of the fragment with the disorder annotations from DisProt, IDEAL, and MobiDB as obtained above; a residue was considered disordered if it was annotated as such in any of these three databases. Fragments comprising at least 20% disordered residues were considered as IDR fragments whereas the others were considered as non-IDR fragments. This cutoff balances an over-representation of ordered residues while ensuring enough training data. This is also consistent with proteome-wide analyses which indicate that proteins with IDRs have 13–28% disordered residues. Specifically, annotations based on experiments indicate a 17% disorder fraction ^35^.

Subsequently, a set of binary complexes was constructed from each PDB entry. Each binary complex consisted of an IDR fragment paired with another IDR or non-IDR fragment. Binary complexes comprising solely of intra-chain fragments or non-IDR fragments were excluded (Fig 1). This resulted in a total of 2369712 binary complexes.

#### Creating merged binary complexes

We then created contact maps from these binary complexes. A contact map is a binary matrix of zeros (non-contacts) and ones (contacts); two residues are in contact if the distance between their Cα atoms is less than 8□ . We merged the contact maps across different binary complexes of a sequence fragment pair, to create “merged contact maps” (Fig 1). This allows us to account for contacts from all available complexes. Further, a merged contact map is less sparse and therefore easier to learn compared to an ensemble of contact maps. It also mitigates the bias caused by having different numbers of structures for different sequence fragment pairs.

We first grouped the binary complexes across all the PDB entries by the UniProt ID of the constituent proteins in the sequence fragment pair, resulting in 14599 UniProt ID pairs/groups. For computational efficiency, all binary complexes with no contacts were eliminated. The remaining binary complexes in a group were sorted based on the UniProt start residue positions of the protein fragments in the pair and further partitioned into sets. Each set comprised binary complexes containing overlapping sequences (more than one residue overlap) for both fragments such that the sequence length for each fragment, merged across the overlapping sequences, does not exceed the maximum fragment length (100 residues). Contact maps for all binary complexes in a set were aggregated using a logical OR operation to form “merged contact maps”. A sequence fragment pair with the corresponding merged contact map comprises a merged binary complex. We obtained 24442 merged binary complexes. Finally, we filtered merged binary complexes for which the contact density was very high (>5%) or very low (<0.5%), resulting in 6018 merged binary complexes (Fig 1).

#### Creating the training set and OOD test set

We created an out-of-distribution (OOD) test set that is sequence non-redundant with our training dataset and the PDB70 dataset used for training AlphaFold. We first removed merged binary complexes whose sequence fragments were derived from a PDB chain in PDB70. We then obtained a non-redundant set of sequences by clustering our dataset of sequence fragments and the PDB70 cluster representatives using MMSeqs2 ^88^. MMSeqs2 was used at a 20% sequence identity threshold and cluster mode 1. Since the PDB70 dataset is already clustered at 40% sequence similarity, we consider only the representative cluster members for sequence identity comparisons.

For the OOD test set, we selected clusters without any chain from PDB70 that are singleton (only one cluster member *i.e.* self) or doublet (two member sequences, both belonging to the same merged binary complex) clusters. This resulted in 53 merged binary complexes for the OOD test set. We removed an OOD test set entry for which AF2 predictions could not be obtained, resulting in 52 OOD test set complexes. The remaining dataset was split into a training (train), development (dev), and an in-distribution (ID) test set in the ratio 0.9:0.05:0.05, resulting in 5362 train, 298 dev, and 297 ID test set merged complexes.

### Disobind Model Architecture and Training

#### Notations

Here, we define the notation used in this paper.

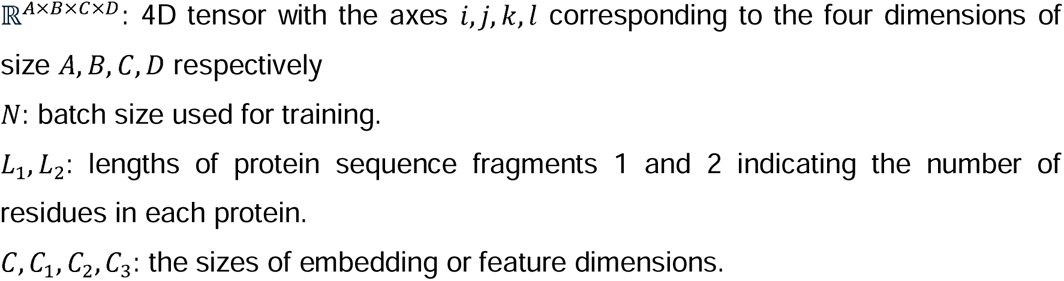

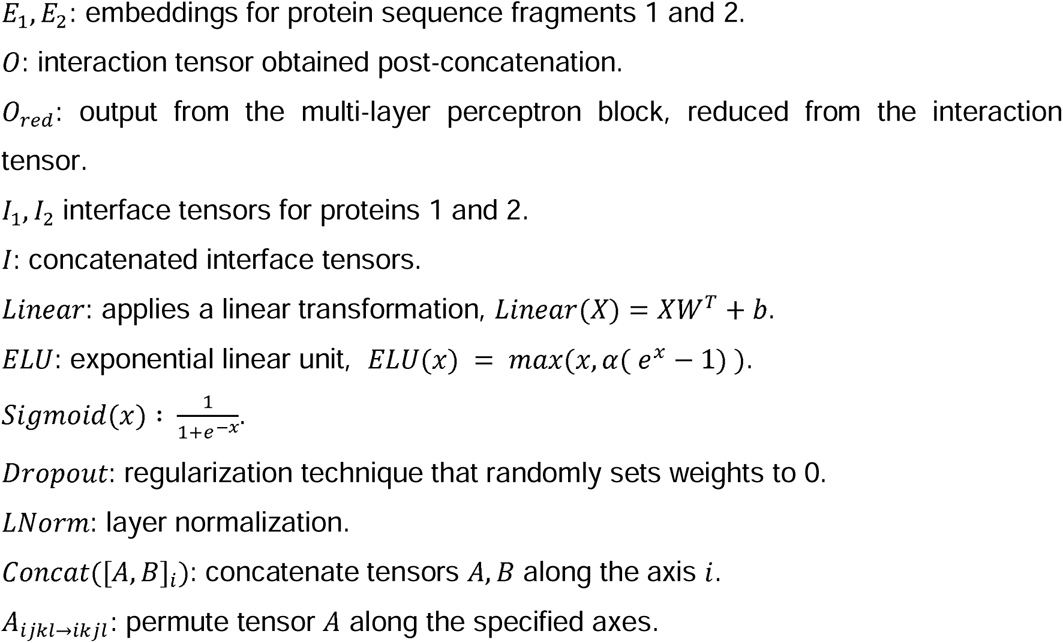

#### Inputs and outputs for training

We train six separate models, corresponding to six prediction tasks. These tasks correspond to two types of predictions: a) inter-protein contact map and b) interface residue predictions at three coarse-grained (CG) resolutions (1, 5, and 10 sequence-contiguous residues). Coarse-graining at resolution 1 is equivalent to no coarsegraining, *i.e*., residue-level predictions. The input to the model is the embedding for the two sequence fragments from the ProtT5 pLM 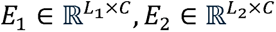 ^44^. The outputs for the two types of predictions respectively are a) a binary contact map, 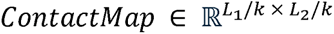 and b) a binary interface residue vector, *Interface* ε 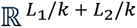 at a particular CG resolution, where k ε [1, 5, 10] represents the CG resolution.

For coarse-grained predictions, the input embeddings were coarse-grained using an AvgPool1d whereas the corresponding contact maps were coarse-grained using a MaxPool2d; the kernel size and stride were set to the CG resolution in both cases.

The target vectors for interface residue predictions were obtained by reducing the rows and columns of the corresponding contact maps by an OR operation. The input embeddings, output contact maps, and interface residue vectors were zero-padded to the maximum length (100 residues).

#### Model architecture

Disobind comprises a projection block, an interface block, a multi-layer perceptron (MLP) block, and an output block (Fig 2). The models for contact map and interface residue prediction employ a contact map block and interface block respectively.

#### Projection block

The input embeddings *E*_1_, *E*_2_ are projected to a lower dimension, *C*_1_ using a linear layer.

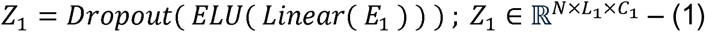

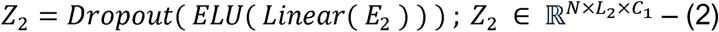

#### Interaction block

The model captures complementarity between the projected embeddings *Z*_1_, *Z*_2_ by concatenating the outer product and the absolute value of the outer difference between the projected embeddings along the feature dimension.

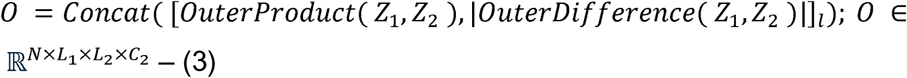

#### Interface block

To capture the interface residues for both proteins *l*, we compute row-wise and column-wise averages followed by concatenating along the length.

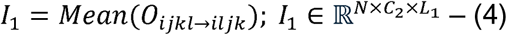

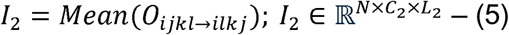

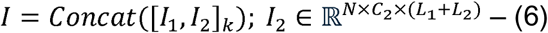

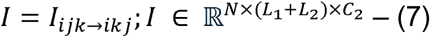

#### Multi-layer perceptron (MLP) block

The feature dimension of the interaction tensor *O* is progressively decreased using a multilayer perceptron (MLP) to obtain *O_red_*.

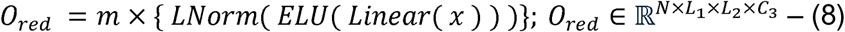

We used a 3-layer (*m* = 3) MLP for contact map prediction whereas it was not used for interface residue prediction (m = 0).

#### Output block

It comprises a *Liner* layer followed by a *Sigmoid* activation.

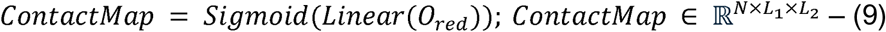

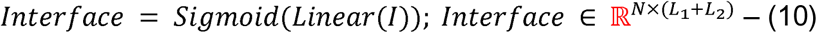

#### Training

The models were trained using PyTorch version 2.0.1 and CUDA version 11.8, using an NVIDIA A6000 GPU. The model was trained using the Singularity Enhanced (SE) loss function ^37^, a modified form of the binary cross-entropy (BCE) loss used for class-imbalanced training, with α = 0.9, β < 3 and AdamW (amsgrad variant) optimizer with a weight decay of 0.05. We used an exponential learning rate scheduler for slowly decaying the base learning rate.

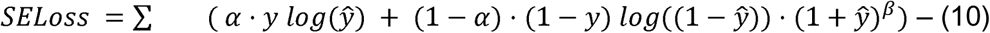

A binary mask was used to exclude the padding from the loss calculation. The train and dev set loss were monitored to check for over/under-fitting (Fig S1).

#### Evaluating predictions

The Disobind predicted score was converted to a binary output using a threshold of 0.5. The model performance was evaluated on the train, dev, and ID test set using recall, precision, and F1-score. All metrics were calculated using the torchmetrics library.

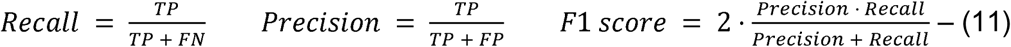

#### Hyperparameter tuning and ablations

Here, we explore the effect of select hyperparameters on the model performances in the dev and ID test sets. Hyperparameters were tuned for the contact map and interface prediction at CG 1 and minimally modified for the CG 5 and CG 10 models. The final set of hyperparameters is provided (Table S1).

##### Projection dimension

We tuned the projection dimension *C_1_* for the projection block (Table S2).

##### Number of layers in the MLP

*Linear* layers in the MLP were used to either upsample (upsampling layers, US) or downsample (downsampling layers, DS) the feature dimension by a factor of 2. We tuned the number of US (DS) layers. Additionally, we removed the MLP for interface residue prediction (Table S3).

##### α, β for SE loss

We tuned the α and β parameters for SE loss; these parameters weigh the contribution of the contact and non-contact elements to the loss (Table S4).

##### pLM embeddings

We compared the model performance using embeddings from ProtT5 (*C* = 1024), ProstT5 (*C* = 1024), ProSE (*C* = 6165), and ProtBERT ^44,54,55^ (*C* = 1024).

Further, we compared two types of embeddings: global and local embeddings (Fig S4, Table S11). Local embeddings were obtained using the sequence of the fragment as input to the pLM. In contrast, global embeddings were obtained by using the complete UniProt sequence as input to the pLM and extracting fragment embedding from it.

### Assessment

#### Random baseline

To compute random baseline predictions, we predicted random contacts and interface residues based on their frequencies in the training set. For the prediction tasks involving coarse-graining, these frequencies were determined from the coarse-grained contact maps and coarse-grained interface residues in the training set.

#### AlphaFold predictions on the OOD test set

We used a local AF2 installation and the AF3 webserver for obtaining OOD test set predictions ^18,27^. AF2 was run with the default arguments as specified in their GitHub repository. Due to the failure of AF2 to relax several of the OOD test set entries we switch off the Amber relaxation step and consider the unrelaxed models for further evaluation. Additionally, we removed an OOD test set entry for which we were unable to obtain the AF2 prediction. We considered the best models from AF2 (based on the ipTM+pTM score) and AF3 (based on the ranking score) for comparison with Disobind. The ipTM score was used to assess the overall model confidence and the pLDDT and PAE metrics assessed the per-residue confidence. pLDDT scores higher than 70 and PAE values lower than 5 are considered as confident ^89^. An ipTM score higher than 0.8 represents a high-confidence prediction whereas an ipTM score lower than 0.6 represents a likely incorrect prediction; predictions with ipTM scores between 0.6-0.8 require further validation ^18,27,45,46^. Given that only five AF2 predictions and none of the AF3 predictions in the OOD test set have an ipTM score greater than 0.8, we initially applied an ipTM cutoff of 0.75 as recommended in benchmarks on AlphaFold2 ^45,46^. Later we relaxed the ipTM cutoffs to 0.4 and 0 (*i.e*., no ipTM cutoff).

To compare the AlphaFold outputs to those from our models, we first derive the binary contact maps from the best-predicted model. We then apply binary masks on this output to ignore low-confidence interactions (pLDDT<70 and PAE>5). Both the pLDDT and PAE masks are applied on the CG 1 contact map obtained from AF2 and AF3 before proceeding further. To apply an ipTM cutoff, we zero the contact maps for AlphaFold predictions with an ipTM lower than the cutoff. The binary contact map was zero-padded to the maximum length (100 residues). The interface residues and coarse-grained predictions were derived from the CG 1 contact map as mentioned previously (See Inputs and Outputs for training).

#### Disobind+AF2 predictions

The Disobind+AF2 predictions were obtained by combining the corresponding predictions from Disobind and AF2 using a logical OR operation. We ignore low-confidence AF2 predictions having a pLDDT less than 70 and a PAE greater than 5. No ipTM cutoff was used.

#### Performance by residue type

Given a protein sequence, disordered regions were identified by combining DisProt, IDEAL, and MobiDB annotations as described in Creating dataset for Disobind section. Residues in these regions were annotated as “disordered residues”, the remaining residues in the sequence were considered “ordered”. Linear interacting peptide (LIP) annotations were obtained from MobiDB as described earlier. We categorized residues into disorder-promoting amino acids (R, P, Q, E, G, S, A, K), aromatic amino acids (F, Y, W), hydrophobic amino acids (A, V, L, I, P, M, F, W), and polar amino acids (S, T, C, N, Q, Y, D, E, K, R, H) ^90,91^. For LIPs, we evaluated contacts between residues in LIPs and any residue on the partner protein. For all other residue types, we evaluated contacts between residues of the same type. To assess the predictions on specific residue types, we applied a binary mask on the predictions and the target to ignore interface residues (contacts) not involving the relevant residues (residue pairs). We only considered confident predictions from AF2 having pLDDT scores higher than 70 and PAE values lower than 5. No ipTM cutoff was used.

#### Comparison with interface predictors for IDRs

We selected state-of-the-art interface predictors including AIUPred, MORFchibi, and DeepDISOBind for comparison to Disobind on the OOD test set ^25,49,50^. These methods have been shown to be comparable to the state-of-the-art in CAID2 ^48^. They provide partner-independent interface residue predictions. For comparison to Disobind, we assess their interface residue predictions on the IDR sequence of the binary complexes in the OOD set. AIUPred predictions were obtained using programmatic access as mentioned in their GitHub repository. For MORFchibi and DeepDISOBind, predictions were obtained using their respective webservers, using defaults. DeepDISOBind provides binary interface predictions. AIUPred predictions are converted to binary interfaces using a 0.5 threshold as specified in the paper ^49^. For MORFchibi, we use a 0.77 threshold for interface residues as recommended; interface residue regions shorter than four residues are eliminated ^50^.

### Quantification and statistical analysis

Not applicable

### Additional resources

Not applicable

## Notes

https://github.com/isblab/disobind

### Competing Interest Statement

The authors have declared no competing interest.

### Summary of Updates

Version updated based on revisions recommended by the journal (adding some data and adding more text in the introduction)

