## Supplementary material for "Disobind: a sequence-based, partner-dependent contact map and interface residue predictor for intrinsically disordered regions"

### Figures

**Figure S1. Loss plots for training Disobind.** We use the singularity-enhanced (SE) loss function for training Disobind. The loss per epoch is calculated for the train and dev set and monitored for under/over-fitting. Related to Methods: Training.

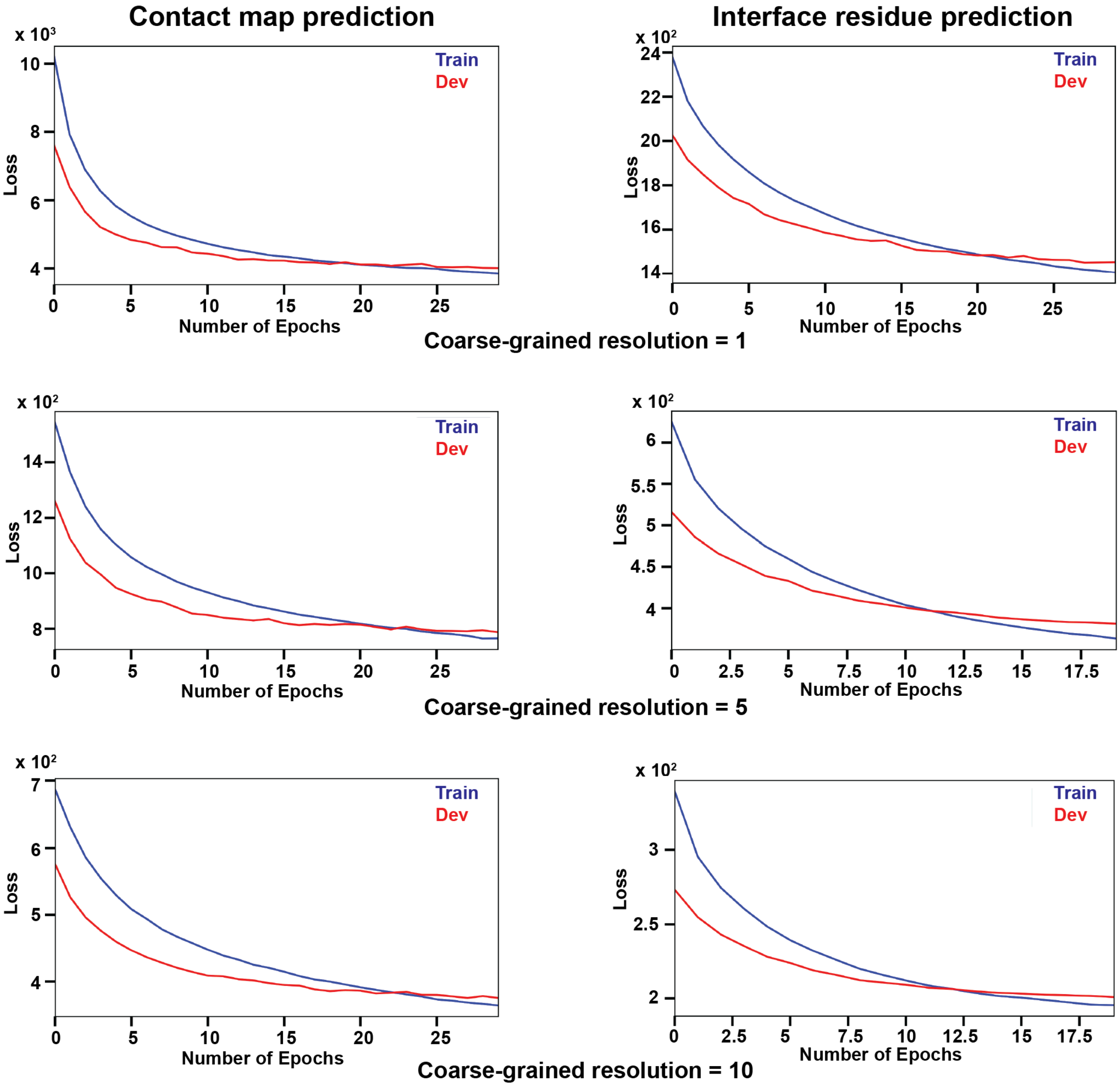

**Figure S2. Plot of sparsity vs OOD set performance.** A plot for the sparsity in the training set vs the model performance measured by the F1 score on the OOD set. We calculate sparsity as the percentage of one minus the fraction of contacts or interface residues in the entire training dataset. Here (C1, C5, C10), (I1, I5, I10) refer to the contact map and interface residue prediction, respectively, at coarse-grained resolutions 1, 5, 10. Related to Figure 3.

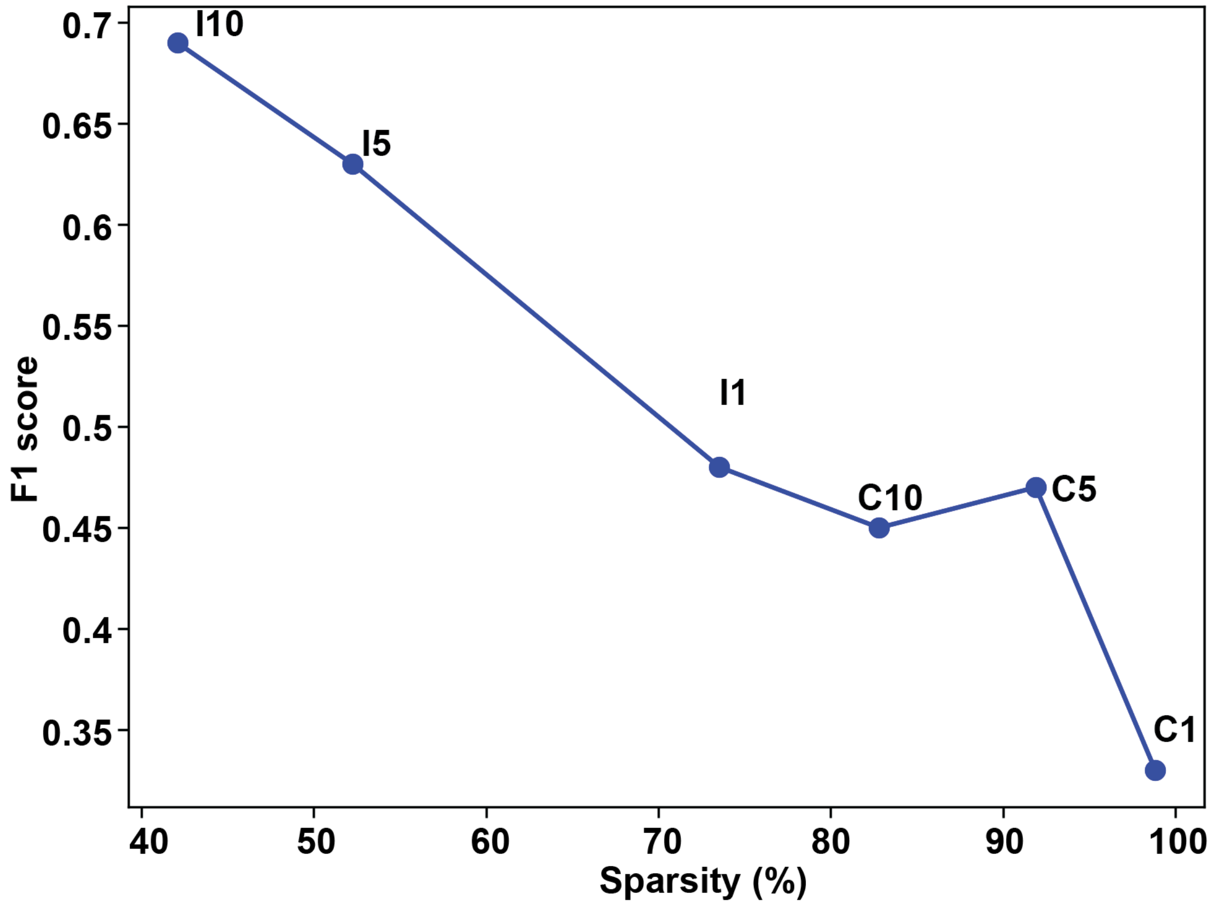

**Figure S3. Comparison of AF2 and AF3 model confidence.** Comparing the model confidence as measured by the ipTM for AF2 and AF3 predictions across all entries in the OOD set. Related to Results: Comparison to AlphaFold2 and AlphaFold3 - AF2 performs better than AF3.

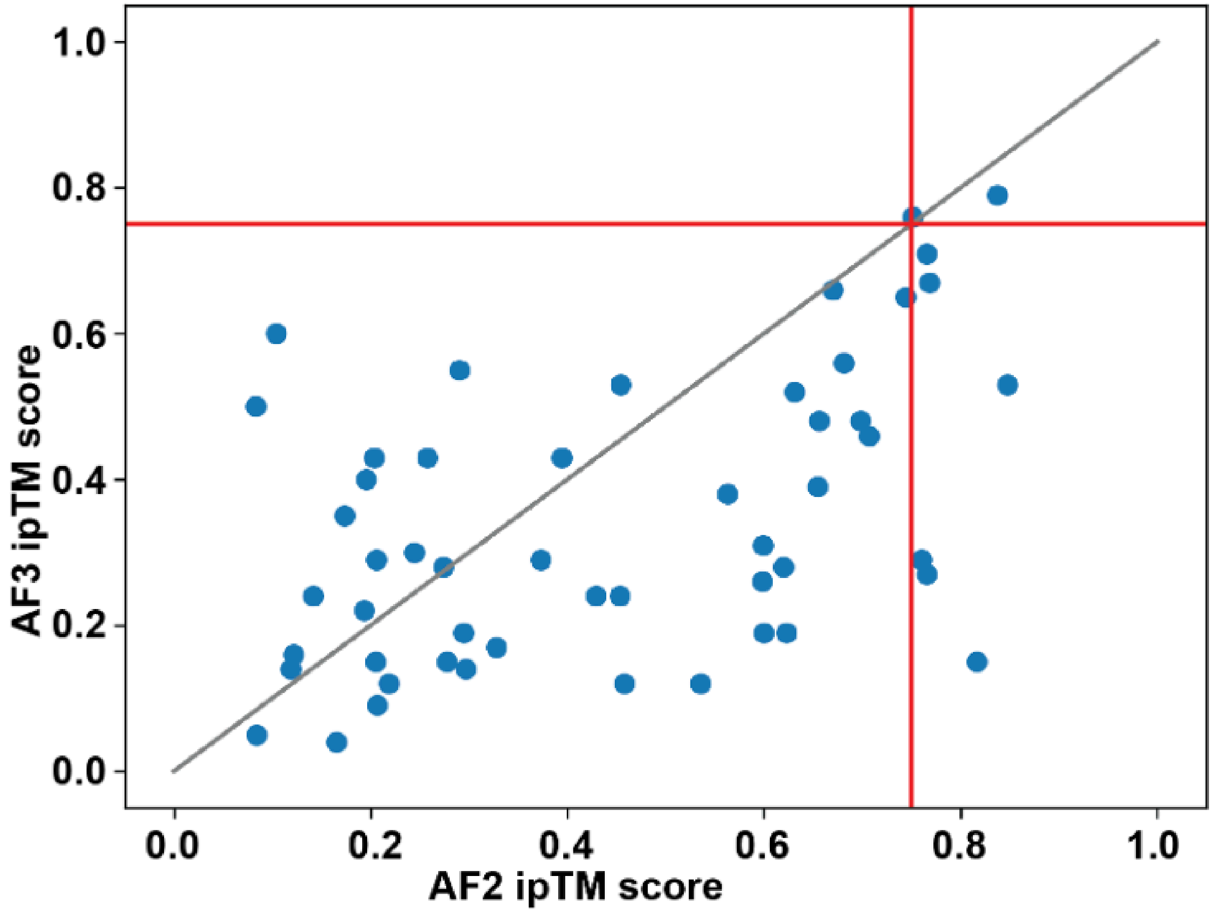

**Figure S4: Global and local sequence embeddings.** We obtain global and local embeddings for the input protein sequence fragments. For global embedding, the complete UniProt protein sequence corresponding to the sequence fragment (dashed blue box) is used as input to the pLM from which the sequence fragment embedding is extracted. For local embeddings, the sequence fragment (dashed blue box) is used as input to the pLM to obtain the embedding. Global embeddings provide the context of the flanking sequence of IDRs that may modulate their binding. Related to Results: Input Embeddings.

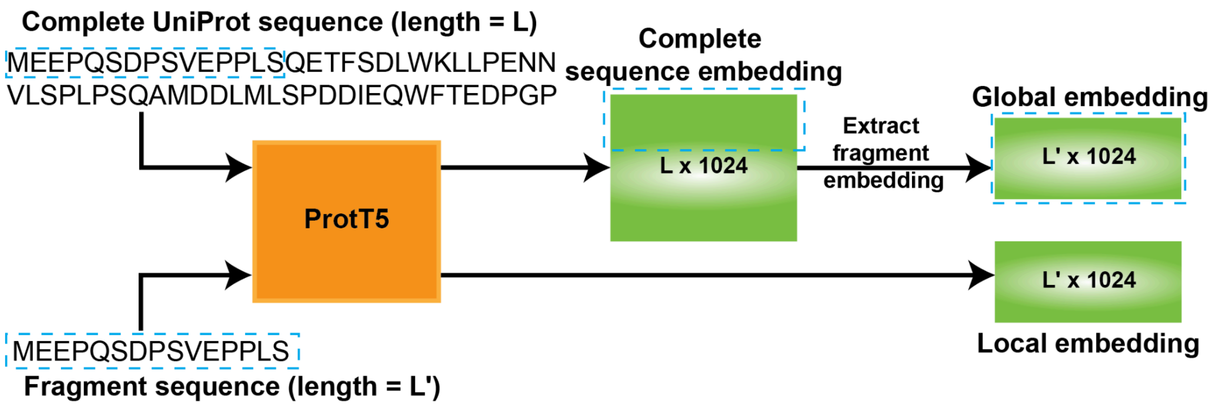

**Figure S5. Visualizing Disobind+AF2 predictions for IDR complexes.** Visualizing the Disobind+AF2 predicted and target interface residues for biologically important IDR complexes. Many of these are involved in phase separation and remain disordered upon binding (DDR). Examples from MobiDB, PED, BMRB, and LLPSdb were selected for which NMR structures were available. The protein chains are coloured in blue and red, where the lighter shade represents the chain and the darker shade represents the interface residues for the respective chains. For all NMR structures, the ensemble is represented with reduced transparency in the background, whereas the first model is shown in the foreground. The PDB IDs, protein names, and corresponding UniProt accession, along with the residue numbers for which the predictions were made, are shown for all complexes. Related to Figure 4.

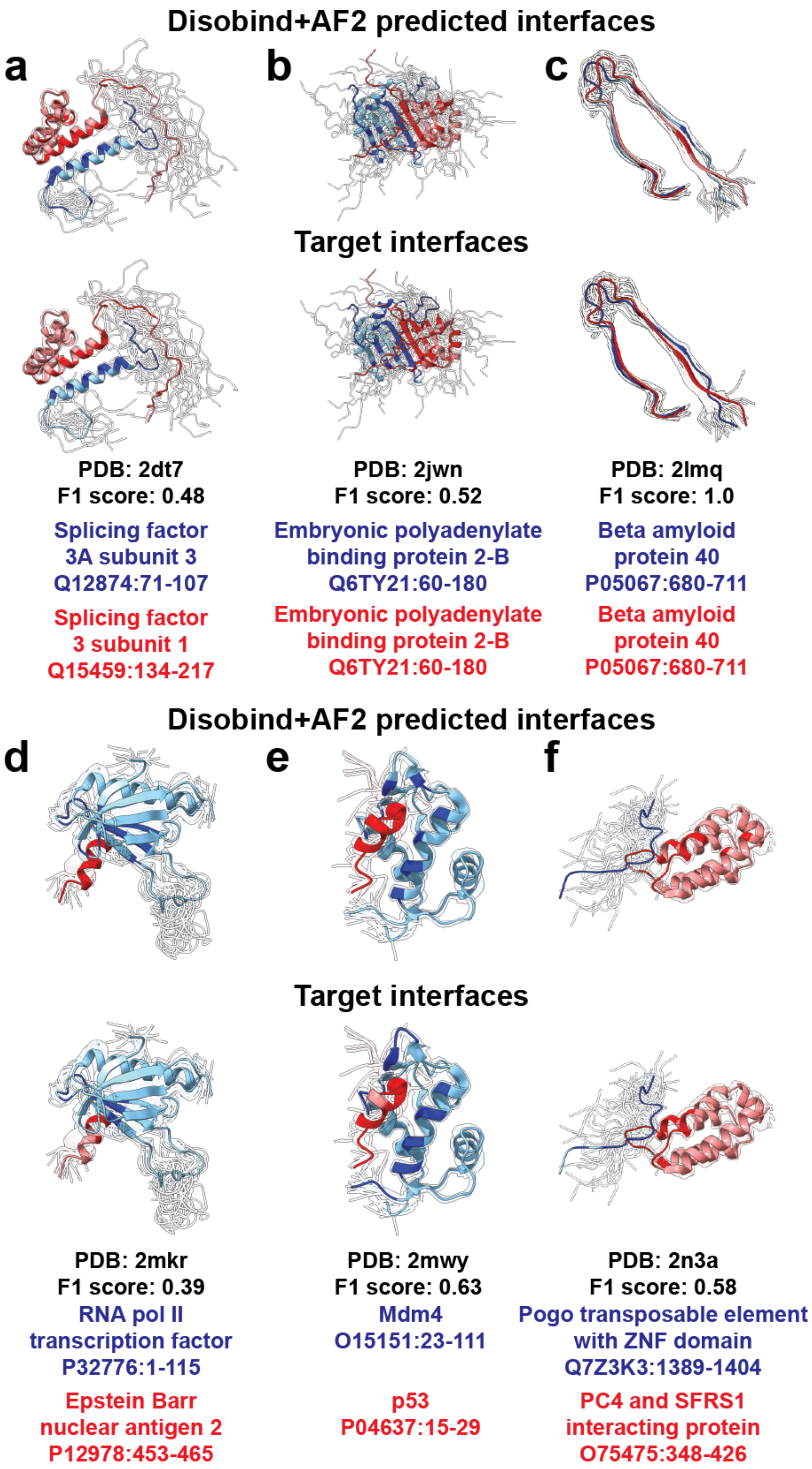

Disobind+AF2 predicted interfaces
a
b
c
Target interfaces
PDB: 2dt7
F1 score: 0.48
Splicing factor
3A subunit 3
Q12874:71-107
Splicing factor
3 subunit 1
Q15459:134-217
PDB: 2jwn
F1 score: 0.52
Embryonic polyadenylate
binding protein 2-B
Q6TY21:60-180
Embryonic polyadenylate
binding protein 2-B
Q6TY21:60-180
PDB: 2lmq
F1 score: 1.0
Beta amyloid
protein 40
P05067:680-711
Beta amyloid
protein 40
P05067:680-711
Disobind+AF2 predicted interfaces
d
e
f
Target interfaces
PDB: 2mkr
F1 score: 0.39
RNA pol II
transcription factor
P32776:1-115
Epstein Barr
nuclear antigen 2
P12978:453-465
PDB: 2mwy
F1 score: 0.63
Mdm4
O15151:23-111
p53
P04637:15-29
PDB: 2n3a
F1 score: 0.58
Pogo transposable element
with ZNF domain
Q7Z3K3:1389-1404
PC4 and SFRS1
interacting protein
O75475:348-426

### Supplementary Tables

**Table S2. Tuning the projection dimension.** The input embeddings are projected to a lower dimension (projection dimension) by the projection block. The performance reported here is on the dev set for predictions at coarse-grained (CG) resolution 1. Related to Methods: Training - Hyperparameter tuning and ablations.

| Projection dimension | Recall | Precision | F1-score |
|---|---|---|---|
| | Contact map prediction, CG 1 | | |
| **128** | 0.49 | 0.67 | 0.57 |
| **256** | 0.5 | 0.68 | 0.57 |
| **512** | 0.47 | 0.68 | 0.56 |
| | Interface residue prediction, CG 1 | | |
| **64** | 0.74 | 0.66 | 0.7 |
| **128** | 0.76 | 0.66 | 0.71 |
| **256** | 0.75 | 0.68 | 0.71 |

**Table S3: Tuning the number of layers in MLP.** Upsampling (US) and downsampling (DS) layers were used in the MLP for the contact map and interface residue prediction tasks. The performance reported here is on the dev set for predictions at coarse-grained (CG) resolution 1. Related to Methods: Training - Hyperparameter tuning and ablations.

| No. of hidden layers | Recall | Precision | F1-score |
|---|---|---|---|
| | **Contact map prediction, CG 1** | | |
| **DS 3** | 0.5 | 0.68 | 0.57 |
| **DS 2** | 0.52 | 0.66 | 0.58 |
| **DS 1** | 0.49 | 0.69 | 0.57 |
| **US 1** | 0.5 | 0.67 | 0.57 |
| **US 2** | 0.49 | 0.69 | 0.56 |
| | **Interface residue prediction, CG 1** | | |
| **DS 2** | 0.72 | 0.7 | 0.71 |
| **DS 1** | 0.75 | 0.65 | 0.7 |
| **DS 0** | 0.76 | 0.66 | 0.71 |
| **US 1** | 0.78 | 0.63 | 0.69 |
| **US 2** | 0.77 | 0.63 | 0.69 |

**Table S4: Tuning SE loss parameters α and β.** We tested various combinations of the SE loss parameters α and β for training Disobind. The performance reported here is on the dev set for predictions at coarse-grained (CG) resolution 1. Related to Methods: Training - Hyperparameter tuning and ablations.

| Parameter | Recall | Precision | F1-score |
|---|---|---|---|
| | **Contact map prediction** | | |
| **α=0.8, β=3** | 0.41 | 0.82 | 0.54 |
| **α=0.8, β=2** | 0.48 | 0.71 | 0.57 |
| **α=0.9, β=2** | 0.55 | 0.55 | 0.55 |
| **α=0.9, β=3** | 0.5 | 0.68 | 0.57 |
| | **Interface residue prediction** | | |
| **α=0.8, β=3** | 0.6 | 0.77 | 0.67 |
| **α=0.8, β=2** | 0.74 | 0.66 | 0.7 |
| **α=0.9, β=2** | 0.86 | 0.55 | 0.67 |
| **α=0.9, β=3** | 0.76 | 0.66 | 0.71 |

**Table S5: Disobind performance on the in-distribution test set.** Performance of Disobind on the ID test set for contact map and interface residue prediction for different coarse-grained (CG) resolutions. Related to Results: Evaluating Disobind.

| CG | Recall | Precision | F1-score |
|---|---|---|---|
| | **Contact map prediction** | | |
| **1** | 0.49 | 0.69 | 0.57 |
| **5** | 0.58 | 0.7 | 0.64 |
| **10** | 0.66 | 0.7 | 0.68 |
| | **Interface residue prediction** | | |
| **1** | 0.73 | 0.67 | 0.7 |
| **5** | 0.86 | 0.73 | 0.79 |
| **10** | 0.92 | 0.76 | 0.83 |

**Table S6: Sparsity in the dataset for the contact map and interface residue prediction task for all coarse-grained resolutions.** We calculate the sparsity in our dataset for contact map and interface residue prediction for different coarse-grained (CG) resolutions as the fraction of positives, i.e., contacts or interface residues, in the entire training dataset. Related to Figure 3.

| Task | Coarse Graining (CG) | Fraction of positives |
|---|---|---|
| **Contact map prediction** | 1 | 0.0117 |
| | 5 | 0.081 |
| | 10 | 0.1719 |
| **interface residue prediction** | 1 | 0.2646 |
| | 5 | 0.4774 |
| | 10 | 0.5791 |

**Table S7: Comparing AF3 predictions derived from the structure and from the distogram head.** We compare the performance of AF3 for contact map and interface residue predictions obtained from the predicted structure and the contact probabilities predicted from the distogram head. Evaluation is done on the OOD test set, considering only the confident predictions (pLDDT >= 70 and PAE <= 5) with no ipTM cutoff applied. Related to Results: Comparison to AlphaFold2 and AlphaFold3.

| | **CG** | **Contact maps from AF3 structure** | **Contact maps from AF3 contact probabilities** |
|---|---|---|---|
| **Contact map prediction F1-score** | **1** | 0.13 | 0.14 |
| | **5** | 0.22 | 0.23 |
| | **10** | 0.25 | 0.25 |
| **Interface residue prediction F1-score** | **1** | 0.27 | 0.27 |
| | **5** | 0.34 | 0.33 |
| | **10** | 0.37 | 0.35 |

**Table S8: Comparison of Disobind, AF2, and Disobind+AF2 predictions in disordered regions and ordered regions.** We evaluate the performance of Disobind, AF2, and Disobind+AF2 in predicting contacts and interface residues in disordered and ordered regions considered separately. For AF2 and Disobind+AF2, we only consider the confident predictions (pLDDT >= 70 and PAE <= 5) with no ipTM cutoff applied. Related to Results: Performance by residue type.

| Model | Recall | Precision | F1-score | Recall | Precision | F1-score |
|---|---|---|---|---|---|---|
| | Contact map prediction | | | Interface residue prediction | | |
| | Interactions in disordered regions, CG 1 | | | | | |
| **Disobind** | 0.49 | 0.77 | 0.6 | 0.56 | 0.55 | 0.56 |
| **AF2_0.0** | 0.24 | 0.73 | 0.36 | 0.35 | 0.85 | 0.49 |
| **Disobind+AF2** | 0.5 | 0.73 | 0.6 | 0.62 | 0.55 | 0.58 |
| | Interactions in ordered regions, CG 1 | | | | | |
| **Disobind** | 0.17 | 0.48 | 0.25 | 0.43 | 0.39 | 0.41 |
| **AF2_0.0** | 0.23 | 0.34 | 0.27 | 0.36 | 0.6 | 0.45 |
| **Disobind+AF2** | 0.35 | 0.36 | 0.36 | 0.58 | 0.4 | 0.47 |

**Table S9: Comparison of Disobind, AF2, and Disobind+AF2 on predicting contacts and interface residues for various amino acid types and motifs.** We evaluate the performance of Disobind, AF2, and Disobind+AF2 in predicting contacts and interface residues involving specific amino acid types, including disorder-promoting, aromatic, hydrophobic, and polar amino acids. We also compare the performance in predicting contacts and interface residues involving linear interacting peptides (LIPs) in at least one of the proteins. For AF2 and Disobind+AF2, we only consider the confident predictions (pLDDT >= 70 and PAE <= 5) with no ipTM cutoff applied. Related to Results: Performance by residue type.

| Model | Recall | Precision | F1 score | Recall | Precision | F1 score |
|---|---|---|---|---|---|---|
| | Contact map prediction | | | Interface residue prediction | | |
| | Disorder-promoting residues, CG 1 | | | | | |
| **Disobind** | 0.25 | 0.66 | 0.36 | 0.47 | 0.45 | 0.46 |
| **AF2_0.0** | 0.2 | 0.56 | 0.3 | 0.34 | 0.7 | 0.46 |
| **Disobind+AF2** | 0.33 | 0.55 | 0.41 | 0.56 | 0.45 | 0.5 |
| | Aromatic residues, CG 1 | | | | | |
| **Disobind** | 0.19 | 1 | 0.32 | 0.41 | 0.37 | 0.39 |
| **AF2_0.0** | 0.22 | 0.58 | 0.32 | 0.32 | 0.6 | 0.41 |
| **Disobind+AF2** | 0.25 | 0.62 | 0.36 | 0.56 | 0.38 | 0.45 |
| | Hydrophobic residues, CG 1 | | | | | |
| **Disobind** | 0.33 | 0.69 | 0.44 | 0.53 | 0.47 | 0.5 |
| **AF2_0.0** | 0.28 | 0.42 | 0.34 | 0.39 | 0.67 | 0.49 |
| **Disobind+AF2** | 0.43 | 0.47 | 0.45 | 0.63 | 0.46 | 0.53 |
| | Polar residues, CG 1 | | | | | |
| **Disobind** | 0.16 | 0.57 | 0.25 | 0.48 | 0.47 | 0.48 |
| **AF2_0.0** | 0.17 | 0.52 | 0.26 | 0.34 | 0.72 | 0.46 |
| **Disobind+AF2** | 0.27 | 0.51 | 0.35 | 0.59 | 0.49 | 0.53 |

| | Linear Interacting Peptides (LIPs), CG 1 | | | | | |
|---|---|---|---|---|---|---|
| **Disobind** | 0.3 | 0.81 | 0.43 | 0.58 | 0.58 | 0.58 |
| **AF2_0.0** | 0.26 | 0.69 | 0.37 | 0.41 | 0.91 | 0.57 |
| **Disobind+AF2** | 0.41 | 0.7 | 0.52 | 0.64 | 0.59 | 0.62 |

**Table S10: Performance by the type of complexes.** We evaluate the performance of Disobind, AF2, and Disobind+AF2 on OOD test set complexes containing long IDRs, as well as DOR and DDR complexes. For AF2 and Disobind+AF2, we only consider the confident predictions (pLDDT >= 70 and PAE <= 5) with no ipTM cutoff applied. Related to Results: Comparison to AlphaFold2 and AlphaFold3 - Combining Disobind and AlphaFold2 predictions. Related to Results: Comparison to AlphaFold2 and AlphaFold3 - Combining Disobind and AlphaFold2 predictions.

| **Model** | **Recall** | **Precision** | **F1 score** | **Recall** | **Precision** | **F1 score** |
|---|---|---|---|---|---|---|
| | **Contact map prediction, CG1** | | | **Interface residue prediction, CG1** | | |
| | **Long IDR-containing complexes** | | | | | |
| **Disobind** | 0.32 | 0.73 | 0.44 | 0.58 | 0.49 | 0.53 |
| **AF2** | 0.3 | 0.83 | 0.44 | 0.36 | 0.9 | 0.51 |
| **Disobind+AF2** | 0.44 | 0.76 | 0.56 | 0.67 | 0.52 | 0.59 |

**Table S11: Comparison of model performance across multiple pLMs.** We compare global and local embeddings obtained from ProtT5, ProstT5, ProSE, and ProtBERT for both the contact map and interface prediction tasks on the ID test set. Related to Results: Input Embeddings.

| pLM | Embedding Type | Recall | Precision | F1 score |
|---|---|---|---|---|
| | | **Contact map prediction** | | |
| **ProtT5** | **global** | 0.49 | 0.69 | **0.57** |
| | **local** | 0.41 | 0.72 | 0.52 |
| **ProstT5** | **global** | 0.48 | 0.7 | 0.57 |
| | **local** | 0.41 | 0.74 | 0.52 |
| **ProSE** | **global** | 0.47 | 0.68 | 0.55 |
| | **local** | 0.45 | 0.7 | 0.54 |
| **ProtBERT** | **global** | 0.27 | 0.63 | 0.37 |
| | **local** | 0.17 | 0.62 | 0.27 |
| | | **Interface residue prediction** | | |
| **ProtT5** | **global** | 0.73 | 0.67 | **0.7** |
| | **local** | 0.7 | 0.67 | 0.68 |
| **ProstT5** | **global** | 0.72 | 0.69 | 0.7 |
| | **local** | 0.68 | 0.67 | 0.67 |
| **ProSE** | **global** | 0.82 | 0.58 | 0.68 |
| | **local** | 0.78 | 0.64 | 0.7 |
| **ProtBERT** | **global** | 0.63 | 0.62 | 0.62 |
| | **local** | 0.57 | 0.59 | 0.58 |
